# Competition strategies driving resource partitioning in chitin degrading communities

**DOI:** 10.1101/2024.11.07.622309

**Authors:** Sammy Pontrelli, Ghita Guessous, Julian Trouillon, Aswin Krishna, Terence Hwa, Uwe Sauer

## Abstract

Resource competition strongly shapes microbial community dynamics and function. In polysaccharide-degrading communities, primary degraders secrete monomer-releasing enzymes and exploiters consume released products without investing in enzymes. By investigating competitive strategies between chitin degraders and N-acetylglucosamine (GlcNAc) exploiters, we uncover several interacting factors, including GlcNAc liberation rates, diffusive loss, and resource partitioning between competitors, which determine community viability and growth dynamics at early stages of degradation. Aside from cases involving antibiotic secretion or aggregation on particles to privatize degrading enzymes, exploiters also inhibit degraders by siphoning away limiting GlcNAc flux during early stages of particle degradation, when degraders must overcome diffusive GlcNAc loss to sustain chitinase production. Analogous to the Allee effect in population biology, these effects lead to sensitive dependences on initial species densities for a community to thrive. This study elucidates how metabolic competition at early stages of particle degradation shapes species interactions, resource partitioning, and ultimately community viability.

## Introduction

Resource sharing and competition are important drivers of inter-species dynamics in microbial communities^1–7^. In both laboratory experiments^3,6,8,9^ and complex environments like the mammalian gut^10^, microbes with similar metabolic capabilities often compete for resources leading to species exclusion; yet some communities can sustain metabolically similar species coexisting on the same resources^11–16^. This raises questions about the relative influence of initial community composition, spatial structure, and non-metabolic forms of inhibition on inter-species competition. Thus, to fully understand species interactions and their coexistence in these competitive settings, it is essential to unravel the underlying relationship between competitive strategies and the various mechanisms of resource partitioning.

Competition for publicly available resources is particularly prevalent in polysaccharide degrading communities, which have important implications for human health^17^, biotechnology^18^, and biogeochemical cycling^19^. These communities are characterized by the presence of specialized degraders that release hydrolytic enzymes extracellularly to break down polysaccharides into transportable monomers and oligomers^20,21^. The extracellular degradation of polysaccharides generates publicly available mono- and oligomers that can be utilized not only by the degraders but also by exploiters that cannot produce their own hydrolytic enzymes^21–24^. These competitive dynamics between exploiters and degraders are frequent and affect population dynamics and polysaccharide degradation rates^3,25–27^. This is because due to the presence of exploiters, degraders face diminished returns from their investment in hydrolytic enzymes. Although degraders can increase enzyme expression to enhance carbon flow^28,29^, the deployment of such a strategy is limited by their intake of the consumable monomers and oligomers siphoned away by the exploiters. To enhance monomer access in spatially structured environments, degraders and exploiters can aggregate on polysaccharide surfaces^30–33^ or rapidly disperse to new nutrient patches^31,32,34,35^. To prevent diffusive loss, they may employ membrane-bound hydrolytic enzymes or selfish oligomer uptake^22,36 37,38^. The key question revolves around the potential advantages and costs of different strategies employed by competing species.

Here, we investigate the competitive strategies of degraders and exploiters in a model marine chitin-degrading community. The insoluble form of chitin, the most abundant polysaccharide in the ocean^39^, allows us to explore the role of microbial attachment and aggregation in the competitive dynamics, which has implications for carbon cycling in marine ecosystems and other microbial systems that degrade insoluble polysaccharides^20,40,41^. To identify instances of direct competition for the chitin monomer N-acetylglucosamine (GlcNAc), we screened pairwise cocultures of an 18 bacteria chitin-degrading community (**Table S1**)^12,41^, previously shown to recapitulate population dynamics of chitin-degrading consortia^3^. Out of these cocultures, we characterized five pairs in detail, to identify key factors that influence competition between degraders and exploiters and their growth dynamics. In some cases, we found that antibiotic production enabled degraders to protect themselves while in other cases, colony aggregation on particles helped fend off exploiters. Besides these direct competitive mechanisms, we also uncovered how metabolic competition shapes community dynamics. In particular, initial stages of particle degradation proved critical, when degraders are challenged by resource limitations and diffusive GlcNAc loss, hindering chitinase production necessary for community growth. Exploiters exacerbate these challenges by siphoning precious GlcNAc, delaying or even completely preventing coculture growth. The results reveal sensitive dependences of community viability on initial conditions and early degradation dynamics as a hallmark of resource competition.

## Results

### Screening for altered chitin-degrading phenotypes in pairwise cocultures reveals species-specific interactions

To identify strategies of how degraders and exploiters compete for GlcNAc as it is liberated from chitin, we used a model marine chitin-degrading community that consisted of bacteria previously categorized into three functional guilds based on their chitin degradation abilities^3^: “Degraders” express chitinases to release monomeric GlcNAc, “exploiters” do not express chitinases and depend on released GlcNAc for their growth, while “scavengers” cannot utilize GlcNAc or chitin and rely on other metabolites released by degraders or exploiters. We paired one of five exploiters or scavengers with one of four degraders (20 combinations each) and followed the growth of each coculture at 1:1 initial cell ratio (**Fig. S1, S2**, respectively, with selected combinations shown in **Fig. 1A**). Each species was seeded at a density of 10^7^ cells/mL (OD_600_ of about 0.01, **Table S2**). The impact on degrader growth was assessed by quantifying changes in lag time (**Fig. 1B, Table S3**), defined as the time required for each culture to surpass an OD_600_ of 0.25. Out of the 40 cocultures studied, 19 showed inhibitory interactions (with most lag times increasing by at least a day), and only two exhibited positive interactions (red and blue symbols, respectively, **Fig. 1B**). Most inhibitory interactions (13 out of 19) were caused by exploiters, consistent with the hypothesis that many of these phenotypes are due to competition for GlcNAc. The six cases of degrader inhibition through scavengers (**Fig. S2**) suggest that inhibition for reasons beyond GlcNAc competition can also occur.

**Figure 1.**
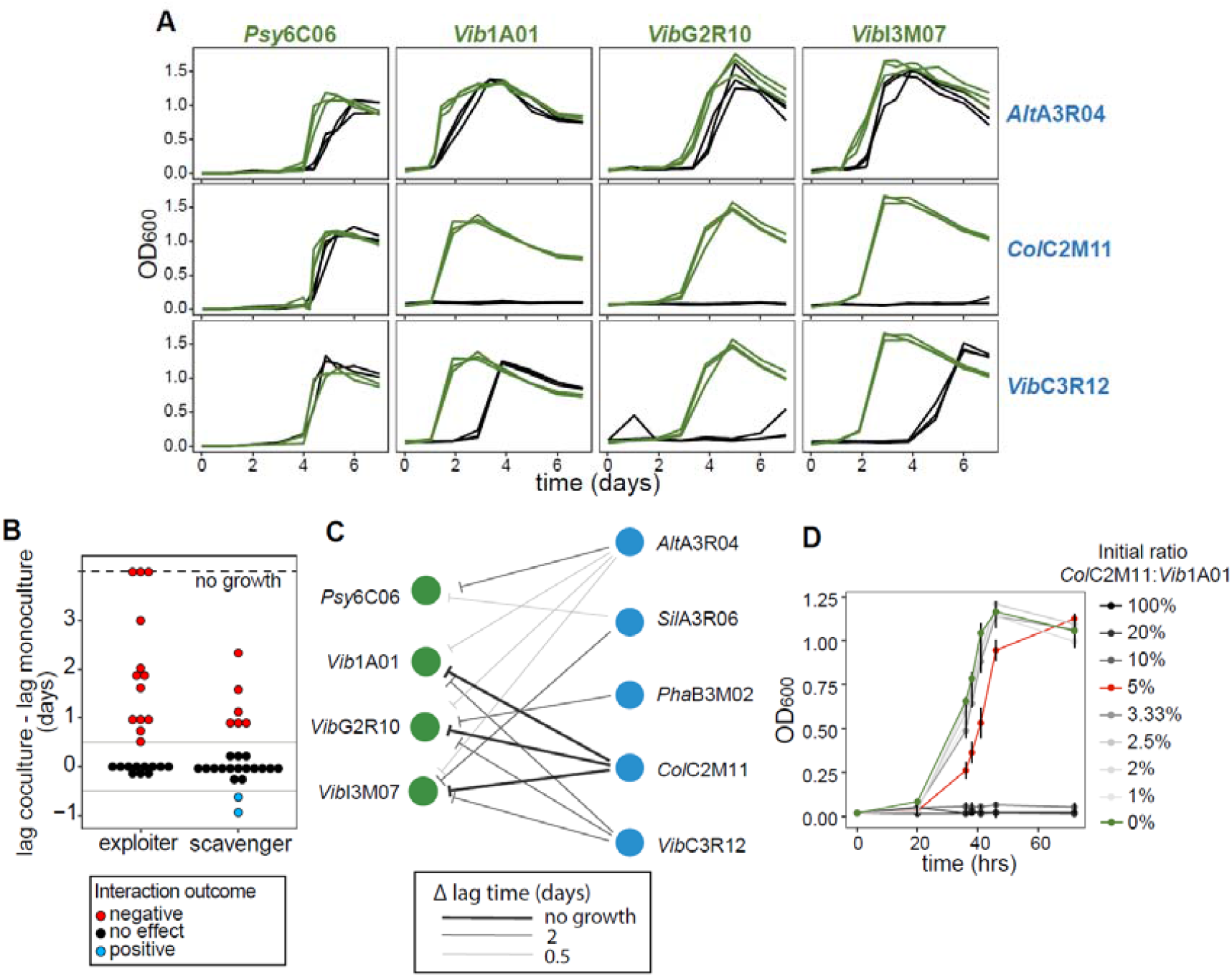
Growth of selected pairwise cocultures during chitin degradation. A) Growth curves of degrader monocultures (green) and degrader-exploiter cocultures (black). B) Change in lag time (time to surpass OD_600_ 0.25), calculated by subtracting the lag time of cocultures from the lag time of monocultures. Red dots represent an inhibition phenotype, where exploiters or scavengers confer a relative increase in lag time greater than 12 hours. Blue dots represent a positive interaction, where the exploiters or scavengers decreases the lag time by more than 12 hours. C) Exploiter (blue) induced lag times of degraders (green). Arrow width represents the increase in lag time induced by the exploiter in coculture. D) Growth on colloidal chitin of *Vib*1A01 with *Col*C2M11 at different inoculation ratios, where *Vib*1A01 is 1×10^7^ cells/mL. Triplicate growth experiments are shown, where error bars are standard deviation of the mean. The remaining pairwise coculture growth curves are in Fig. S1, S2.

In this work, we focus on the interactions between degraders and exploiters (**Fig. 1C**) due to their prevalence. Only *Alt*A3R04 moderately inhibited the growth of all degraders, while the other four exploiters inhibited some degraders strongly but not others. For instance, *Col*C2M11 and *Vib*C3R12 did not inhibit *Psy*6C06, *SilA*3R06 inhibited only *Psy*6C06 and *Vib*I3M07, and PhaB3M02 inhibited only *Vib*G2R10. ColC2M11 was the only exploiter capable of completely inhibiting growth (thick arrows in **Fig. 1C**; see also growth curves in **Fig. 1A**). However, the extent of inhibition was dependent on the initial inoculation density. For example, the complete inhibitory effect of *Col*C2M11 on *Vib*1A01 vanished when *Col*C2M11 was seeded at 20-fold or lower cell density (red curve, **Fig. 1D**). An influence of degrader preculture conditions was ruled out by inoculating cocultures with *Vib*1A01 grown in rich and chitin minimal medium, demonstrating nearly identical inhibition in both cases (**Fig. S3**). Generally, whether the *Vib*1A01:*Col*C2M11 co-culture grows or not seems to depend on the ratio of the initial inoculants (**Fig. S4**). This suggests that species-specific inhibition is influenced by different competitive traits and may be condition dependent.

### Testing for toxin secretion and proximity requirement of inhibitory interactions

To differentiate inhibition caused by competition for GlcNAc from other forms of inhibition, we selected the most inhibitory exploiters *Col*C2M11, *Vib*C3R12, and *Alt*A3R04 that caused 10 of 13 inhibitory interactions (**Fig. 1A, S1**) and tested their effect on degraders. *Col*C2M11 and *Vib*C3R12 were paired with the emerging model chitin degrader *Vib*1A01^3,41–43^, and *Alt*A3R03 was paired with *Psy*6C06 that exhibited the longest increase in lag time compared to the other degraders.

To test whether growth inhibition was due to toxic compound secretion by the exploiter, the three degraders were grown on colloidal chitin in mono- and coculture with their paired exploiters until early stationary phase. *Col*C2M11 was inoculated at a 20-fold lower density than *Vib*1A01 since this allowed some growth of the coculture (**Fig. 1D**). Cell- and chitin-free supernatants were obtained by filtration and diluted 50% with fresh colloidal chitin medium. We determined the difference in lag time when degraders were grown in these diluted supernatants of cocultures and their own monocultures. If the exploiter secretes a toxin, we expect the degrader to exhibit a longer lag phase in the coculture supernatant. Only *Psy*6C06 showed a growth delay in the coculture supernatant (**Fig. 2A, S5**), indicating that *Alt*A3R04 secretes an inhibitory compound. Filtration of the supernatant through a 10 kDa molecular weight cutoff filter confirmed that *Alt*A3R04 appears to release a low molecular weight inhibitor (**Fig. 2A, S5**).

**Figure 2.**
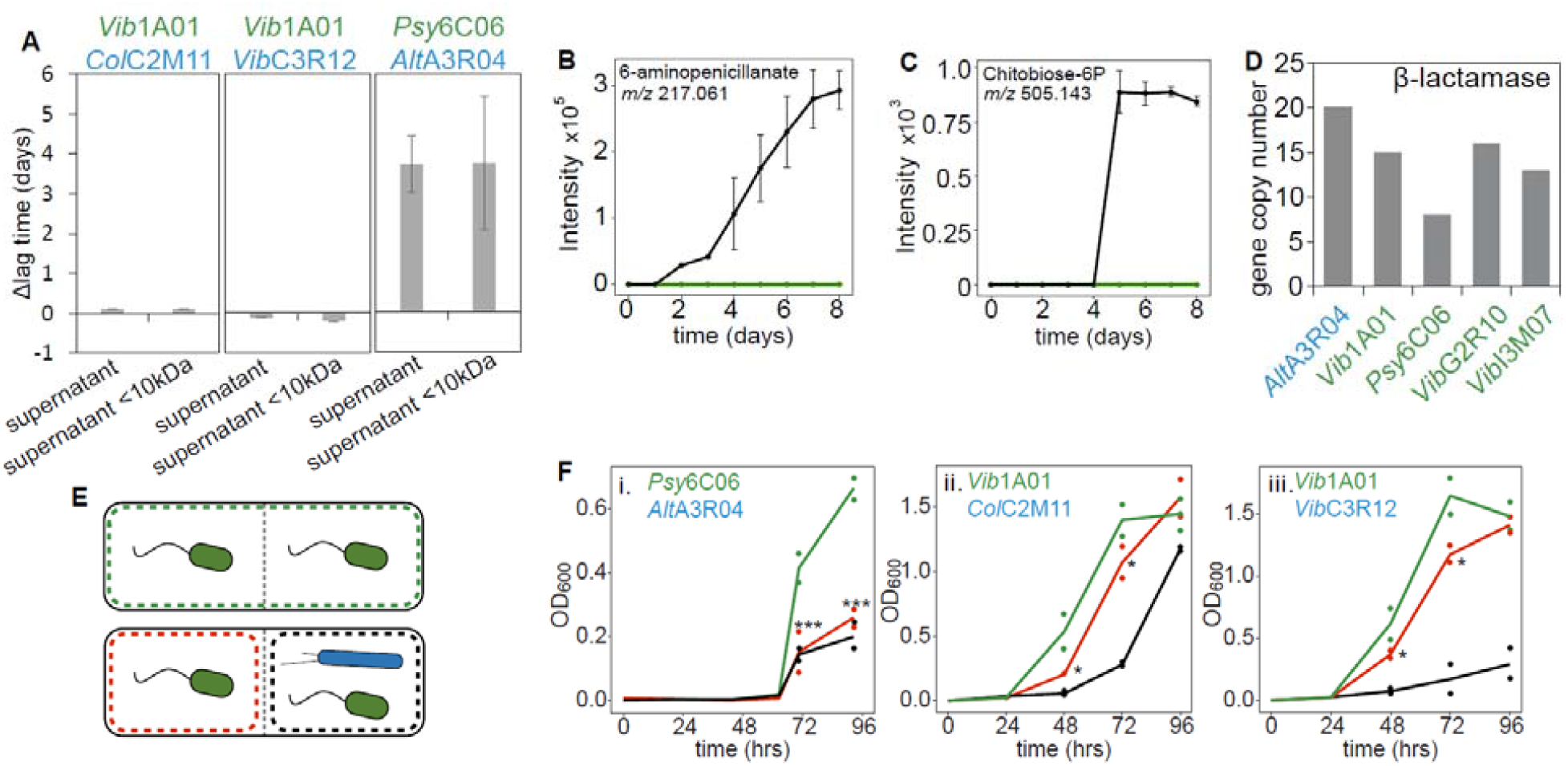
Degrader (green) inhibition through exploiter (blue) toxin secretion or direct contact. A) Change of degrader lag time in monoculture on colloidal chitin diluted with 50% supernatant from degrader:exploiter cocultures or degrader monocultures after growth on chitin. Change in lag time to reach an OD_600_ of 0.25 is defined as the lag time in the supernatant from monoculture minus the lag time in supernatant from the coculture. B) Metabolomics measurement of 6-aminopenecillanate in *Psy*6C06:*Alt*A3R04 coculture (black) and *Psy*6C06 monoculture (green) on colloidal chitin. C) Concentration of chitobiose-6-phosphate in *Psy*6C06:*Alt*A3R04 coculture (black) and *Psy*6C06 monoculture (green). D) Copy number of annotated β-lactamases in exploiter *Alt*A3R04 and four degraders. E) Setup of membrane-separated coculture chambers with degraders (green) and exploiter (blue). F) Growth in membrane-separated growth chambers: degrader monocultures on both sides (green lines) and degrader monoculture (red lines) separated from degrader:exploiter coculture (black lines). Coculture experiments were performed in duplicate, cells were inoculated at 1×10^7^ cells/mL. Stars represent significant changes (*p* < 0.05, one-tailed t-test) when comparing the degrader monocultures with or without the exploiter inoculated on the other side of the membrane (i.e., comparison of red and green lines only). All other experiments were performed in triplicate and all error bars represent the standard deviation of these replicates.

Untargeted liquid chromatography quadrupole time of flight mass spectrometry (LC-QTOF-MS) revealed accumulation of 41 putatively annotated metabolites in coculture of *Psy*6C06 and *Alt*A3R04 compared to *Psy*6C06 monoculture on colloidal chitin (**Supplemental Dataset 1**). Of interest is 6-aminopenicillanate (6-APA, **Fig. 2B**), a precursor or degradation product of β-lactam antibiotics^44^, which also possesses antibiotic properties by disrupting cell wall biosynthesis^45,46^ leading to cell lysis^47^ in Gram-negative bacteria. *Alt*A3R04, possesses three genes encoding penicillin amidases, which can produce 6-APA from β-lactam compounds^48^. Although β-lactam antibiotics like Penicillin G were undetectable (data not shown), accumulation of phosphorylated chitobiose implied toxin-dependent cell lysis and release of intracellular chitin breakdown products from *Psy*6C06 (**Fig. 2C**), because *Alt*A3R04 is unable to take up chitobiose^3^, the GlcNAc dimer. We hypothesize that *Alt*A3R04 produces a β-lactam antibiotic that inhibits *Psy*6C06 and possibly other degraders in coculture (**Fig. 1A**). Interestingly, all organisms in this study carry multiple β-lactamase genes (**Fig. 2D, S6, Supplemental Dataset 2**), with *Alt*A3R04 having the highest copy number (20), presumably to protect itself against any β-lactams it secretes, and *Psy*6C06 is the degrader having the lowest number (8).

Since the cell-free supernatant experiments did not show any toxin-dependent inhibition through *Col*C2M11 and *Vib*C3R12 **(Fig. 2A, S5)**, we tested whether spatial proximity was a requirement of the inhibitory dynamics. To do this, we used a device that separates two culture chambers with a 0.1 μm pore size membrane, allowing the passage of metabolites or secreted toxins, but not of cells^49^. With colloidal chitin as a sole carbon source, we inoculated each of the three degraders in both membrane-separated chambers and their corresponding exploiter in only one chamber **(Fig. 2E)**. The extent of inhibitory effects resulting from the coculture would be apparent when comparing growth in devices that contain only the degrader in each chamber (top cartoon, **Fig. 2E**) to devices where one chamber contains the degrader only and the other one, a degrader-exploiter coculture (bottom cartoon, **Fig. 2E**). Interactions requiring the co-localization of the two species would exclusively occur in the co-culture chamber (black box in **Fig. 2E**) while interactions mediated by diffusible molecules would only be visible in the mono-culture chamber (red box in **Fig. 2E**). Using this setup with colloidal chitin as a sole carbon source, we tested three degraders and their corresponding exploiter.

First, addition of *Alt*A3R04 affected the growth of *Psy*6C06 strongly and similarly in both chambers, which reinforces the above finding that *Psy*6C06 is inhibited across distance by a diffusible small molecule (**Fig. 2F,i**). For *Col*C2M11 and *Vib*C3R12, some inhibition occurred in the adjacent degrader-only chamber (compare the red and green curves, **Fig. 2F,ii-iii**). This allows us to rule out inhibitory processes that depend exclusively on cell-to-cell contact. The data on *Col*C2M11 and *Vib*C3R12 suggest an intriguing scenario: while physical proximity is important for the inhibition of the degrader *Vib*1A01, small molecules that diffused across the coculture membrane also play a role. Since spent medium from stationary phase did not have an inhibitory effect (**Fig. 2A**), these small molecules could be nutrients liberated during growth, e.g., monomers or oligomers of GlcNAc, suggestive of metabolic competition.

### GlcNAc uptake kinetics and metabolic competition

For chitin cultures to progress through the lag phase, eventual growth sustaining both species depends on the ability of the degrader to access sufficient resources for chitinase production. Since the inhibitory effects of *Col*C2M11 and *Vib*C3R12 on *Vib*1A01 likely involve some aspects of metabolic exchange, we next examined the ability of the exploiters to compete with *Vib*1A01 for GlcNAc, which depends on substrate affinity and maximum growth rate^50–52^. Although *Vib*1A01 grows twice as fast at higher GlcNAc concentrations than *Col*C2M11 (**Fig. 3A**), the exploiters may have uptake kinetics that allow them to outcompete the degrader at low concentrations. This regime is reflective of the initial phase of particle degradation, where the nutrient concentration is expected to be very low due to the low flux of nutrient generation.

**Figure 3:**
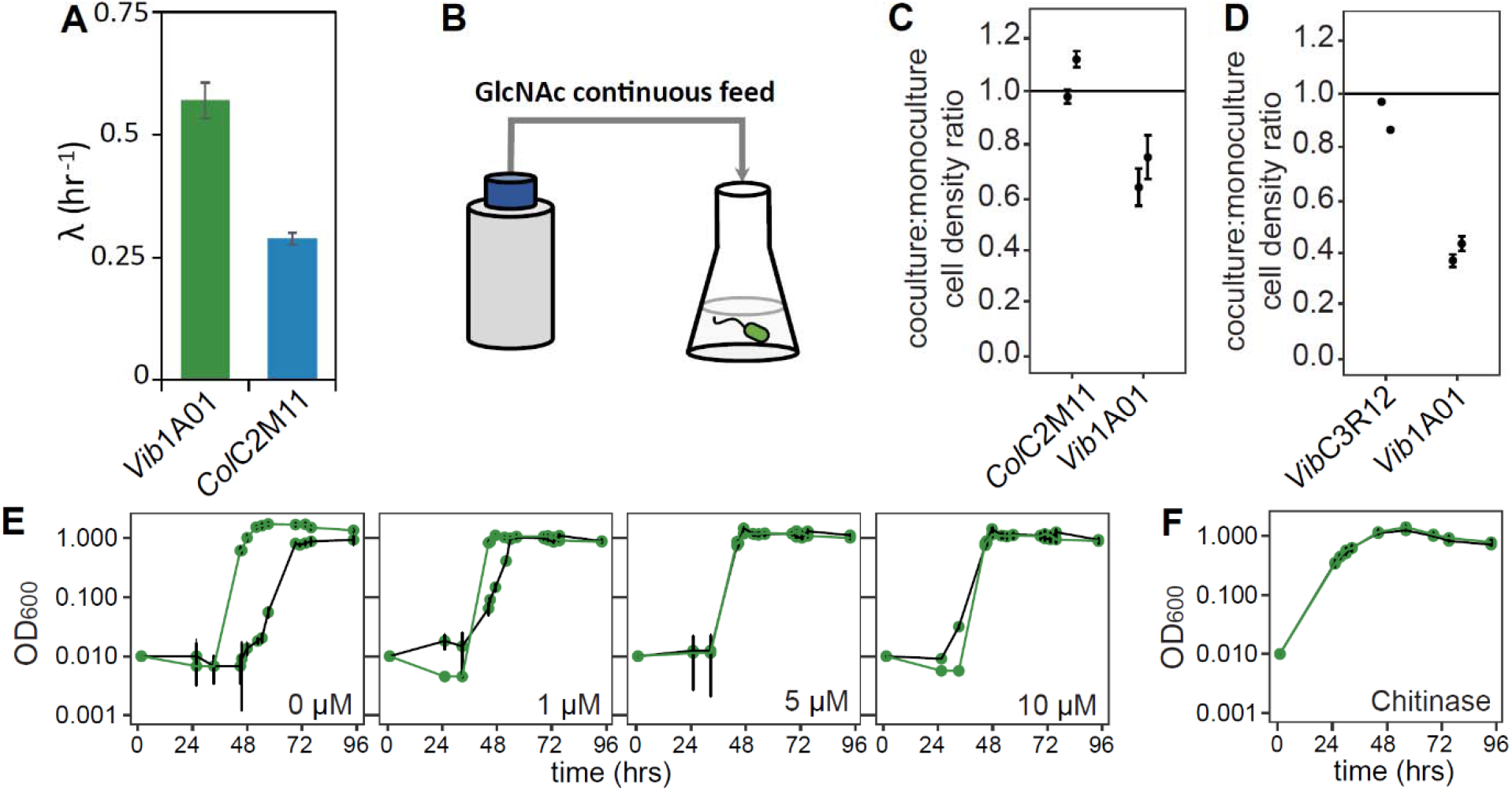
Maximum growth rates and species abundance in GlcNAc-limited fed-batch cultures. A) Maximum growth rates in 20 mM GlcNAc minimal medium in batch culture. B) Schematic of a GlcNAc-limited fed-batch reactor. GlcNAc is continuously dripped into a culture of the inoculated species that initially contains no carbon source. C-D)Relative species abundance in GlcNAc-limited fed-batch cocultures compared to monoculture for C) *Vib*1A01 with *Col*C2M11 and D) *Vib*1A01 with *Vib*C3R12. Each species was inoculated at 1×10^7^ cells/mL and cell density was determined after 72 hr. Horizontal black lines are a visual guide, which correspond to a ratio of 1, indicating no change. E) Growth of *Vib*1A01 monoculture (green) and coculture (black) with *Col*C2M11 with various initially spiked in, non-growth supporting concentrations of GlcNAc. F) Growth of *Vib*1A01 monoculture (green) and coculture (black) with *Col*C2M11 upon addition of purified chitinase at inoculation. Error bars represent standard deviation of the mean from triplicate batch cultures and triplicate qPCR measurements of duplicate fed-batch cultures.

To test the hypothesis of inferior degrader uptake kinetics, we competed the degrader *Vib*1A01 with each of the two exploiters (*Col*C2M11 and *Vib*C3R12) in a fed-batch reactor. Bioreactor cultures were inoculated at 10^7^ cells/ml for each species, with the only carbon source being the continuously supplied GlcNAc to mimic gradual GlcNAc release during chitin hydrolysis by the degrader (**Fig. 3B**). The rate of GlcNAc feeding (15 μM/hr initially and 7.5 μM/hr towards the end of the 72h growth period) was adjusted to mimic the rate of GlcNAc formation observed in bulk *Vib*1A01 monocultures between 0-24 hours (∼24 μM/hr at OD_600_ 0.02). Under such nutrient limitation, species specialized to consume scarce nutrients are expected to outcompete others.

We assessed the abundance of each species in the coculture after 72 h by qPCR, normalizing species abundance in the coculture to the abundance of the corresponding mono-culture in the fed-batch reactor with the same GlcNAc feeding rate and the same initial inoculation (**Fig. 3C, 3D**). Assuming that the GlcNAC yield of these species is similar, the result allows us to compare the growth rate of the two species in low GlcNAc concentrations. Instead of the complete dominance by the exploiters, we find that the abundance of *Vib*1A01 dropped only moderately, to ∼2/3 that of *Col*C2M11 and ∼1/2 that of *Vib*C3R12 after 72 h of growth.

These abundance data can be used to estimate quantitatively the ratio of the growth rates of the two competing species in fed-batch growth (**Supplemental Note 1**). The less than 2-fold reduction in *Vib*1A01 abundance found after 72h of competition leads to ∼20% difference in the growth rate of the competing species in the low nutrient regime. This moderate difference does not come close to accounting for the strong inhibitory effects exerted by the exploiters (**Fig. 1A**), particularly, the complete inhibition of co-culture growth when 10% (but not 5%) of *Col*C2M11 is added to the initial inoculant of *Vib*1A01 (**Fig. 1D**).

Despite the metabolic similarity of the degrader and exploiters at low GlcNAc concentrations, we find that co-cultures grown on colloidal chitin are sensitive to the addition of small amounts of GlcNAc: the addition of 1-5 μM of GlcNAc at the inoculation of the co-culture eliminated the lag between the mono and co-culture (**Fig. 3E**), suggesting a benefit of these low initial concentrations to *Vib*1A01. This supports the previous conclusion that *Vib*1A01 has a comparable uptake efficiency as the exploiters at low GlcNAc concentration. Moreover, this suggests that the inhibitory effect exerted by the exploiters only needs to involve a small change in the flow of GlcNAc. As ultimately the growth of the coculture involves the synthesis of chitinases by *Vib*1A01, we verified that the addition of a small amount of chitinases to the co-culture alleviated not only inhibition by exploiters but eliminated lag altogether (**Fig. 3F**). Together, these data suggest that both the lag and the lengthening of the lag exerted by the exploiters have to do with them exploiting small amounts of GlcNAc generated at early stage of coculture growth.

### Role of spatial structure in competitive dynamics

An important aspect brought out by the data in **Fig. 2F** is the effect of spatial proximity on inhibition by exploiters: the chamber containing the degrader alone showed reduced growth in the coculture chamber (bottom panel, **Fig. 2E**) compared to the monoculture chamber (top panel, **Fig. 2E**). In addition, the fact that as little as 5 μM GlcNAc eliminated the inhibition (**Fig. 3E**) indicates that this proximity effect is not due to disruptive effects exerted by the exploiters, such as the degradation of chitinases. While *Col*C2M11 and *Vib*C3R12 prevented or delayed the growth of *Vib*1A01 through proximity in cocultures, *Psy*6C06 cocultures were unaffected (**Fig. 1A**). To confirm that *Psy*6C06 was indeed not affected by the exploiters, the final abundances of each species were determined after complete chitin degradation. Indeed, *Psy*6C06’s abundance was comparable in both mono- and cocultures (**Fig. 4A**). To test whether *Psy*6C06 could potentially outcompete *Col*C2M11 and *Vib*C3R12 due to favorable uptake kinetics, GlcNAc-limited fed-batch cultures were again conducted. *Psy*6C06 does not outcompete the exploiters (**Fig. 4BC**). In both cases, the abundance of the exploiters remained similar in mono- and cocultures, while *Psy*6C06 reaches a moderately lower abundance in coculture with *Vib*C3R12, similar to the fed-batch competition results between the exploiters and *Vib*1A01 (**Fig. 3CD**). This suggests that *Psy*6C06 does not possess superior GlcNAc uptake affinities compared to the exploiters.

**Figure 4:**
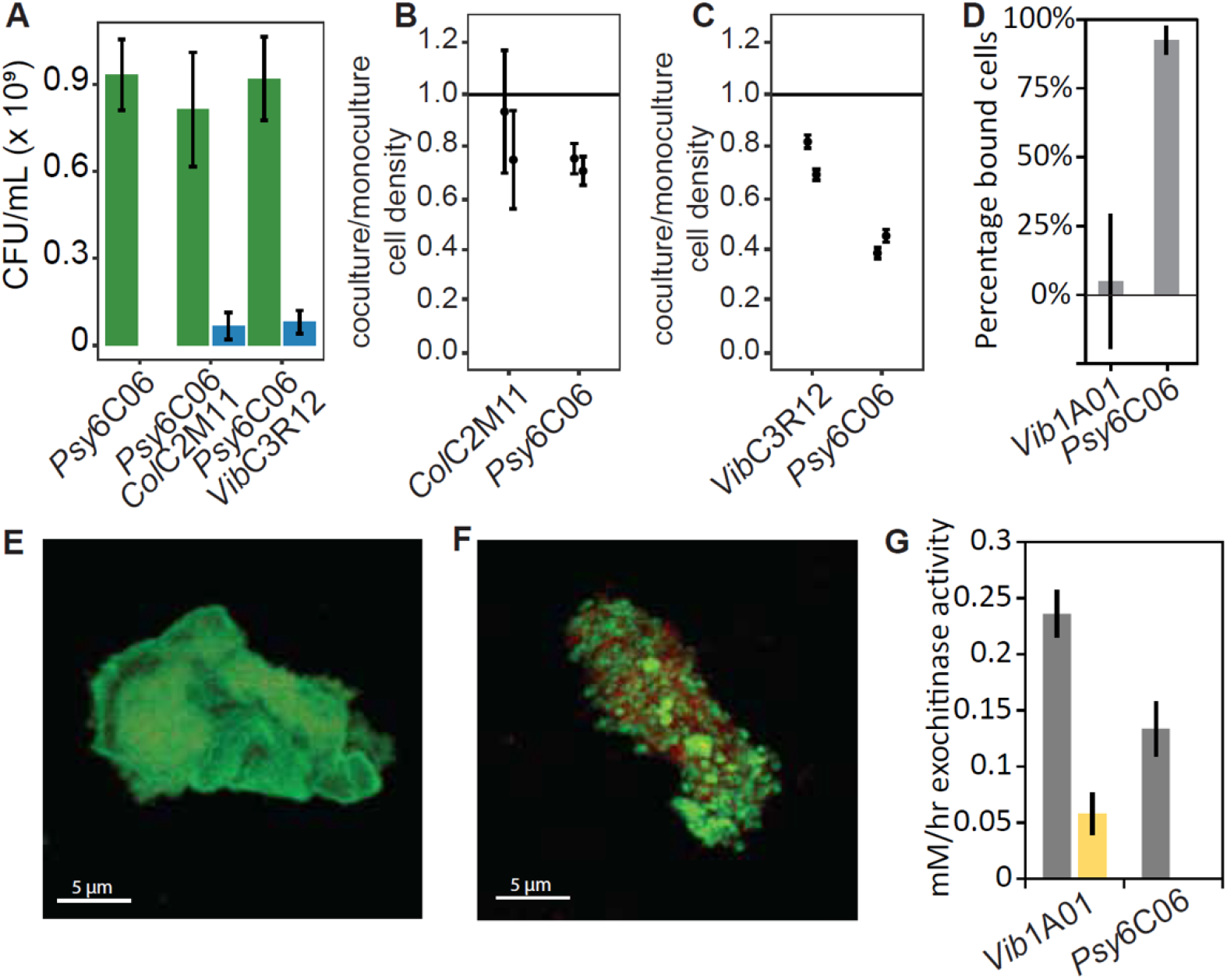
Particle binding compensates for Monod uptake kinetics. A) Early stationary phase colony forming units of *Psy*6C06 (green) and exploiters (blue) in mono- or cocultures on colloidal chitin. B-C) Relative species abundance in GlcNAc-limited fed-batch cocultures compared to monoculture for B) *Psy*6C06 with *Col*C2M11 and C) *Psy*6C06 with *Vib*C3R12. Each species was inoculated at 1×10^7^ cells/mL and cell density was determined after 72 hr. D) Percentage of qPCR determined cells bound to colloidal chitin six hours after inoculation of monocultures. E-F) Microscope images of E) *Vib*1A01 and F) *Psy*6C06 growing on colloidal chitin. Red are the cells; green is the chitin. G) Exochitinase activity in the planktonic (yellow) and chitin-bound (grey) fraction of each degrader after 24 hours of growth on colloidal chitin. In all panels, error bars represent standard deviation of the mean from triplicate experiments and triplicate qPCR measurements of duplicate fed-batch cultures.

One possible explanation for *Psy*6C06’s resilience to the exploiters is its ability to bind to the chitin particle surface^30^. This binding allows *Psy*6C06 to privatize its chitinases and localize itself to regions where chitinase and GlcNAc concentrations are highest. To test this hypothesis, we determined the fraction of each degrader that binds to colloidal chitin after six hours. *Psy*6C06 was almost completely bound, while *Vib*1A01 showed no significant binding (**Fig. 4D**). This is consistent with a previous study which found a minor replicating fraction of *Vib*1A01 on the particle surface, with a larger planktonic and non-dividing population^34^. Using microscopy, we tested the binding of *Vib*1A01 and *Psy*6C06 on the particle surface during logarithmic growth of the monoculture. As expected, our images show dense colonies of *Psy*6C06 bound to the chitin surface, unlike *Vib*1A01 which rather appear to be dispersedly bound across the particle surface (**Fig. 4EF, Supplemental video 1 and 2**). We further measured exochitinase activity on the particle and in the planktonic phase for both degrader monocultures at 24 hours (**Fig. 4G**). In both cases the activity was higher on the particle surface, consistent with previous findings that *Vib*1A01 attaches chitinases to chitin particles^34^. These results suggest that *Psy*6C06 gains its competitive advantage by aggregating on the chitin surface. This aggregation allows *Psy*6C06 to privatize its chitinases and localize itself to the site of GlcNAc liberation, thereby increasing its access to resources and potentially enhancing its competitiveness in the coculture environment. This also suggests that the inability of *Vib*1A01 to exclude exploiters on chitin particles may be a key vulnerability that allows it to be outcompeted when enough exploiters are present initially. The co-localization of exploiters on sites where *Vib*1A01 is breaking down chitin possibly accounts for the large inhibitory effect seen in coculture chambers (black curves, **Fig. 2F**).

### A model of metabolic competition recapitulates co-culture sensitivity to initial conditions

The ability of degraders to establish growing cultures on chitin stems from the positive feedback between chitinase production by the degrader and its uptake of the liberated GlcNAc, that in turn fuels further chitinase synthesis. Qualitatively, an inhibitory effect of exploiters on degraders can arise from breaking this positive feedback: by siphoning GlcNAc away from the degraders, the exploiters can undermine the degraders’ ability to synthesize chitinases, preventing particle degradation altogether. However, the puzzling aspect of the data presented above is that the observed inhibitory interaction is very sensitive to the initial exploiter density (**Fig. 1D**). Specifically, cocultures starting with 5% of *Col*C2M11 showed little inhibitory effect (manifested by a moderate lag time), but those starting with 10% of *Col*C2M11 were unable to grow. The origin of this strong sensitivity is unclear, especially given that the uptake efficiencies of *Vib*1A01 and that of the exploiters are comparable (**Fig. 3CD** and **Supplemental Note 1**). The data in **Fig. 2-3** have identified several key ingredients linked to the observed inhibition of *Vib*1A01 by *Col*C2M11 and *Vib*C3R11: metabolic competition for GlcNAc in the early phase of the co-culture, involving spatial proximity of the degrader and the exploiter. Here we construct a metabolic model with minimal ingredients that recapitulates the observed strong sensitivity to inhibition by exploiters.

To be sensitive to small changes in the initial exploiter density, we hypothesize that the degrader-chitin system is poised close to a phase transition where small changes in the initial GlcNAc flux could substantially affect degrader growth. Generally, a strong nonlinearity is needed to amplify small perturbations such as those caused by the presence of exploiters. However, the simple nutrient dynamics depicted in **Fig. 5A**, comprising of Monod growth (green arrow), chitinase synthesis (brown arrow) and Michaelis-Menten enzyme kinetics (purple arrow), contain only linear or even sublinear dynamics and thus lack a source of amplifying nonlinearity.

**Figure 5:**
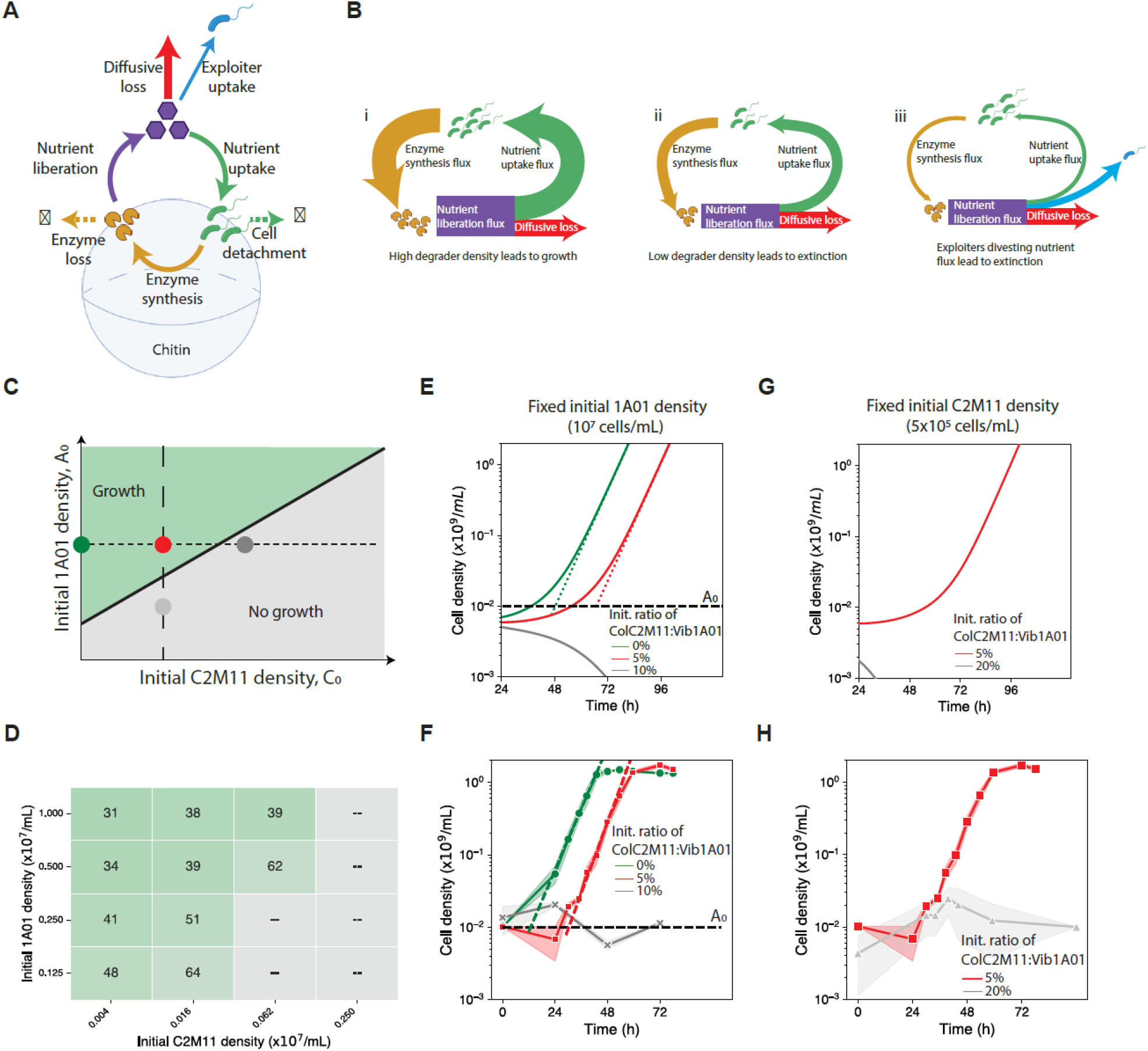
A metabolic model recapitulates phase transition in the growth of the chitin coculture. **A)** Model of chitin-degrader dynamics. Degraders (green cells) synthesize chitinases (yellow pacmans), which in turn generates a GlcNAc liberation flux (purple arrow). The GlcNac molecules (purple hexagons) are taken up by the degraders (green arrow), the exploiters (blue arrow), and also leak away due to diffusion (red arrow). Degraders detach from cells (dashed green arrow) and enzymes turn over (dashed yellow arrow) as empirically determined by Guessous et al^34^. Most parameters in this simple model are experimentally constrained as described in **Supplementary Note 2. B)** Cartoon depicting the partition of the liberated GlcNAc flux in different scenarios: (i) when sufficient amounts of degraders and chitinases are present initially, the GlcNAc liberation flux is large (thick purple box), and cell growth (green arrow) and chitinase production (yellow arrow) is hardly affected by diffusive leakage (red arrow), leading to exponential growth. (ii) At low initial degrader densities/chitinase concentration, the GlcNAc liberation flux is reduced (thin purple box), and most of the GlcNAc liberated is lost to diffusion; this drastically reduces the GlcNAc available for uptake, thus reducing cell growth (as indicated by the thinner green and yellow arrows). (iii) Exploiters can enhance the effect in (ii) by siphoning GlcNAc away (blue arrow) from the degraders and hence from chitinase production, leading to further growth reduction (very thin green and yellow arrows). **C)** An illustrative phase diagram in the space of the initial degrader and exploiter densities. The black line separates the growth and no-growth phases. The green circle represents a monoculture, which is in the growth phase. The red circle indicates 5% exploiter addition, which pushes the system closer to the phase transition. The gray circle indicates 10% exploiter addition, which tips the system to the no-growth phase. **D)** Lag times for chitin coculture growing on various initial degrader and exploiter densities; a snapshot of the coculture OD at 24h after seeding is shown in **Fig. S4**. Lag time is extracted from co-culture growth curves as defined above (OD_600_>0.25). Gray entries correspond to cocultures that did not grow after ∼100h. **E)** Dynamics of the culture according to numerical solutions of the model illustrated in panel A; see **Supplementary Note 2** for model details and also parameters used. The green, red, and gray lines represent the monoculture, and coculture with 5% and 10% initial exploiters, respectively, with the initial degrader at 10^7^ cells/ml (horizontal dotted line, panel C). The monoculture grows exponentially after some lag, with the growth rate (slope of the dashed lines) being 0.16/h. The addition of 5% exploiters (red curve) grows at the same rate, but with the lag increased by ∼20h. The addition of 10% exploiters completely inhibited growth (grey line). **F)** A detailed view of the growth curve for the chitin monoculture of *Vib*1A01 (green) and coculture with *Col*C2M11 (red), with initial cell density being 10^7^ cells/ml and 5×10^5^ cells/ml, respectively. [Grey symbols represent the non-growing coculture with initial density of *Col*C2M11 being 10^6^ cells/ml (10% *Vib*1A01).] The shading represents the standard deviation from three different biological replicates. The population growth rate is 0.18/h for the monoculture and 0.16/h for the coculture with 5% *Col*C2M11, with the latter having ∼18h longer lag. The effect of the exploiters on lag produced by the model is in good agreement with the experiment; but the lag of the monoculture is overestimated in the model due to simplifying approximation made; see **Supplementary Note 2** for discussion. **G)** Dynamics of the cocultures according to the numerical solution of the model, with the initial density of *Col*C2M11 fixed at 5×10^5^ cells/ml, and with varying initial density of *Vib*1A01 (vertical dashed line in panel C): 10^7^ cells/mL (red, same as the red line in panel E), and 2.5×10^6^ cells/mL (gray). The inhibition caused by the exploiter (gray line) is alleviated by higher initial degrader density. **H)** Experimental growth curves of the co-cultures with the same initial conditions as in **G)**. The experimental results are in good agreement with the numerical solution, wherein the gray line exhibits no growth, while the red line shows growth (same as red line in panel F).

Here, we point to a simple physical effect that provides an effective amplifying power to the inhibitory effect of the exploiters on the degraders – the leakage of the liberated GlcNAc away from the degraders by diffusion (red arrow, **Fig 5A**). As described in detail in **Supplemental Note 2**, while the existence of such a diffusive flux is inevitable, it is particularly large for the small colloidal chitin particles studied here, due to the inverse square dependence on surface curvature. Numerically, we estimate that for particles of 10 *μm* radius (**Fig. 4E**), the GlcNAc flux leaked away by diffusion would be comparable to the uptake by *Vib*1A01 cells at a density of 2×10^7^ cells/ml, which is the order of the initial degrader density in our system. This indicates that a density of this order is the minimum necessary to capture a sufficient amount of GlcNAc to get the culture growing (**Fig. 5Bi**,**ii**), i.e., our experimental system is close to the borderline of a phase transition between growth and no growth. It is then plausible that a small reduction in the initial amount of GlcNAc captured by the degraders, resulting from the siphoning of GlcNAc by a small number of exploiters (blue arrow, **Fig. 5A, 5Biii**) could tip the system from growth to no-growth, as illustrated by the phase diagram in **Fig. 5C**. Conversely, a small increase in the initial amount of GlcNAc captured by degraders could ensure growth of the population (**Fig. 3E**). The structure of the growth/no-growth regions in the space of initial degrader and exploiter densities indicated in **Fig. 5C** is echoed qualitatively in **Fig. 5D** by the lag-time data extracted from growth curves collected across many combinations of initial inoculant densities (see **Fig. S4** for a snapshot).

To probe the transition between the two regions more closely, we performed a numerical calculation of the model depicted in **Fig. 5A**; see **Supplementary Note 2**. The model reproduced the basic phenomenon of coculture sensitivity to small changes in the initial exploiter density (**Fig. 5E**). The addition of 5% of exploiters lengthens the monoculture lag by about 20 h (compare the red and green curves), and the addition of 10% of exploiters completely inhibits growth (grey curve), matching the experimental observation (**Fig. 5F, 1D**). In addition to exploring the transition for increasing initial density of *Col*C2M11 at fixed initial density of *Vib*1A01 (horizontal dotted line, **Fig. 5C**), we also probed the transition in the orthogonal direction by changing the initial density of *Vib*1A01 while fixing the initial density of *Col*C2M11 (vertical dashed line, **Fig. 5C**). The same kind of transition is obtained, numerically and experimentally (**Fig. 5G, 5H**, respectively). We note that most parameters used in the numerical calculation are constrained by measurements in this work and from Guessous et al.^34^ for the degrader *Vib*1A01 (listed within Supplementary Note 2), with one experimentally inaccessible parameter (the initial amount of chitinase synthesized by *Vib*1A01 before substantial GlcNAc intake) chosen to place the growth transition at the observed location; see **Supplemental Note 2**. Importantly, the three key ingredients deemed necessary for the occurrence of the abrupt growth transition, the aforementioned diffusive leakage flux, and the turnover of degraders and chitinases on particles, have all been established experimentally for *Vib*1A01^34^. This model quantitatively supports the hypothesis that siphoning of GlcNAc by exploiters during the initial stages can stall particle degradation altogether, and reveals a very sensitive phase transition.

## Discussion

Here, we present molecular and physiological evidence demonstrating how different competition strategies for the chitin monomer GlcNAc contribute to species interactions during polysaccharide degradation. Pairwise coculture experiments revealed widespread species-specific inhibition of degraders through exploiters, each caused by different mechanisms. Notably, the exploiter *Alt*A3R04 was unique in its ability to inhibit all degraders through the secretion of a β-lactam antibiotic. On the other hand, the degrader *Psy*6C06 evaded exploiter competition by binding to chitin particles. This aggregation allows *Psy*6C06 to privatize its chitinases and the resulting degradation products. By doing so, it can concentrate GlcNAc and its oligomers, which in turn enhances per capita growth rates^53^. This strategy protects *Psy*6C06 against exploiters that specialized in growing at low substrate concentrations.

In many cases, the presence of exploiters can completely suppress coculture growth, to the detriment of the exploiters themselves. The interaction dynamics between the degrader *Vib*1A01 and the exploiters *Col*C2M11 and *Vib*C3R12 highlight a fine balance between GlcNAc liberation rates during initial particle degradation, its leakage due to diffusion, and its partitioning between competitors. This interaction is characterized in detail for the degrader *Vib*1A01 and is highlighted by a very sensitive dependence of coculture growth on the initial exploiter density: an addition of 5% of exploiters hardly affected the coculture while 10% of exploiters completely suppressed growth.

An intuitive scenario that may explain this sensitivity to exploiter density is that the exploiters are much better at nutrient competition during the initial phase of chitin degradation, when the concentration of breakdown products is low. However, superior nutrient uptake kinetics by the exploiters would have prevented co-culture growth altogether. Moreover, our study of the fed-batch culture found that the GlcNAc uptake kinetics at low concentrations of GlcNAc were similar for the degrader *Vib*1A01, and the exploiters (**Supplemental Note 1**), invalidating this scenario.

We propose a model based on the fact that *Vib*1A01 detaches rapidly from the chitin particles^34^, such that even in the absence of competition, degraders must accumulate sufficient resources to produce chitinases while overcoming diffusive loss of GlcNAc. In such a scenario, exploiters that compete for GlcNAc further exacerbate GlcNAc limitations caused by diffusive loss, leading to an abrupt growth transition due to small amounts of initial exploiters (**Supplemental Note 2**). As shown in **Fig. 5C-H**, the growth transition driven by changes in the initial density of exploiters is the same as the transition driven by changes in the initial density of degraders. The latter can be viewed as a manifestation of the Allee effect, that the growth of a population (the degraders) requires a critical initial density^53–55^. In this light, metabolic competition can be viewed as a mechanism that increases the critical density of the Allee transition (**Fig. 5C**): the suppression of coculture growth by the exploiters can be understood as the result of the initial density of degraders falling below the now-increased critical density necessary for growth.

Of course, besides the scenario outlined above, there are other plausible scenarios that may account for the observed phenomenon. For example, the degraders can secrete more chitinases to compensate for the low GlcNAc concentration, with a highly sensitive regulation function. However, an advantage of the nutrient competition model is that it does not rely on unknown regulatory processes but is instead built on the system being close to a population-level phase transition point, a property of the system which emerges from known features of the system (particle geometry, Monod kinetics etc.).

Our study highlights the complex dynamics of species-specific interactions and competition, especially during the early stages of chitin degradation, revealing how organisms employ different strategies to gain a competitive edge in nutrient-limited environments. These findings demonstrate that population dynamics are shaped by diverse strategies that different species use to compete and thrive, rather than merely by static partitioning of resources among them. Within our chitin-degrading community, these strategies include toxin secretion, binding to particle surfaces for degraders, as well as siphoning of GlcNAc by exploiters. These findings have implications for other polysaccharide-degrading systems and researchers investigating the impact of resource partitioning on community structure^3,8,11,56–60^. In particular, the very sensitive dependence of community dynamics on the initial species composition, a hallmark of the growth transition described in this study, may occur more broadly in complex polysaccharide-degrading communities, beyond the context of the ocean (for example in the mammalian gut). Understanding the underlying dynamics may be crucial for interpreting community-scale data, which so far has largely been interpreted with a static view, in light of the presence or absence of functions encoded in the genome. In turn, our study emphasizes the importance of dynamics, including initial conditions, for determining the viability of communities, which should be taken into account in interpreting such data and in formulating our understanding of community ecology.

## Methods

### Materials and chemicals

All chemicals were obtained from Sigma-Aldrich unless noted otherwise. Media used are Marine Broth 2216 (Thermo Fisher Scientific, Difco, no. 279100) or MBL minimal medium.

MBL contains 1 mM phosphate dibasic, 1 mM sodium sulfate, and 50 mM Hepes (pH 8.2), and three additional diluted stocks: First, four-fold concentrated seawater salts (NaCl, 80 g/Liter; MgCl2*6H2O, 12 g/Liter; CaCl2*2H2O, 0.6 g/Liter; KCl, 2 g/Liter). Second, 1000 fold concentrated trace minerals (FeSO4*7H2O, 2.1 g/Liter; H3BO3, 30 mg/Liter; MnCl2*4H2O, 100 mg/Liter; CoCl2*6H2O, 190 mg/Liter; NiCl2*6H2O, 24 mg/Liter; CuCl2*2H2O, 2 mg/Liter; ZnSO4*7H2O, 144 mg/Liter; Na2MoO4*2H2O, 36 mg/Liter; NaVO3, 25 mg/Liter; NaWO4*2H2O, 25 mg/Liter; Na2SeO3*5H2O, 6 mg/Liter, dissolved in 20mM HCL). Third, 1000 fold concentrated vitamins (riboflavin, 100 mg/Liter; d-biotin, 30 mg/Liter; thiamine hydrochloride, 100 mg/ liter; l-ascorbic acid, 100 mg/Liter; Ca d-pantothenate, 100 mg/Liter; folate, 100 mg/Liter; nicotinate, 100 mg/Liter; 4-aminobenzoic acid, 100 mg/Liter; pyridoxine HCl, 100 mg/Liter; lipoic acid, 100 mg/Liter; nicotinamide adenine dinucleotide (NAD), 100 mg/Liter; thiamin pyrophosphate, 100 mg/Liter; cyanocobalamin, 10 mg/Liter, dissolved in 10mM MOPS pH 7.2).

### Preparation of colloidal chitin

10 grams of powdered chitin (Sigma-Aldrich, C7170) was dissolved in 100ml of concentrated phosphoric acid (85% by weight) and then placed at 4°C for 48 hours. Roughly 500ml of deionized water was added to this mixture and shaken vigorously until all chitin precipitated. The precipitate was filtered using regenerated cellulose paper (MACHEREY-NAGEL, MN615). Chitin precipitate was then placed in cellulose dialysis tubing (approximately 13kDa, Sigma-Aldrich D9652-100FT) and dialyzed with fresh deionized water daily for three days to remove residual phosphoric acid and oligomers. Following dialysis, the pH was adjusted to 7 with 1M NaOH and homogenized using Bosch SilentMixx Pro blender. The colloidal chitin was sterilized by autoclaving.

### Bacterial strains and culturing

All bacterial species were stored in glycerol stocks at −80°C. Before use, they were streaked onto Marine Broth 2216 plates with 1.5% agar (BD, no. 214010) and placed at room temperature until colonies formed. Overnight precultures were prepared by inoculating a single colony into 2ml of Marine Broth 2216 and placed in a 27°C shaker overnight.

Unless otherwise noted, growth experiments were performed as follows. The medium used is MBL. Cells were inoculated at a density of 1×10^7^ cells/mL. As a carbon source, chitin was supplied at 2 g/L. Cultures were grown in 24 deep well microtiter plates (Kuhner) with 3mL volumes and shaken at 200rpm at 27°C. Growth was monitored with OD_600_ by harvesting 100-200μL of culture for measurement in a Tecan Sunrise plate reader. To prevent suspended chitin from interfering with OD_600_ measurements, the 24 deep well plate was centrifuged for 30 seconds at 1000rcf and the optical density of the supernatant was measured.

### Testing for toxicity in cell-free supernatant

A degrader:exploiter coculture or degrader monoculture was grown on 2g/l MBL in 3mL volumes in a 24 deep well plate in 6 replicates. As soon as the cultures entered into early stationary phase, the supernatant was harvested by centrifuging at 4000rcf for 5 minutes followed by filtration though a 0.22 μM membrane. Large molecules were removed from half of the supernatant using an Amicon Ultra 10kDa cutoff filter before being sterilized again with a 0.22 μM filter. Both supernatants (with and without large molecules removed) were diluted 1:2 with fresh MBL containing 4 g/L chitin, inoculated with the degrader, and growth was measured over time.

### Determining cell abundance with qPCR

Cell abundance in fed-batch reactors were measured using qPCR. Cell cultures were pre-processed as follows: 40μL of culture was added to 10μL of Chelex mastermix (25% wt Chelex 100, 200-400 mesh, and 100mg/mL proteinase from *Aspergillus melle*us Sigma P4032-5G). This mixture was incubated at 56°C for 60 minutes, and 95°C for 10 minutes, and was further diluted 1:10 in nuclease-free water. qPCR was performed using Promega GoTaq qPCR mastermix in 15μL reactions using 6μL of the diluted Chelex reaction as a template. Readings were performed using a QuantStudio 3 (Thermo Scientific) qPCR machine. Genome-specific primers amplified a 75-150bp region on the genome with a measured amplification efficiency of 80-105% (**Table S2**). Cell abundances were determined from the qPCR cycle threshold (Ct) values by comparing Ct values to a standard curve of Ct values measured from samples with known CFU/mL concentrations. All qPCR measurements were performed with two technical replicates that were averaged.

### Chitin binding affinity assay

To determine the fraction of cells that binds to chitin in the first six hours, species were inoculated into MBL containing 2g/l colloidal chitin or no colloidal chitin, in four replicates each. After 6 hours, 500 μL of culture was collected, centrifuged at 1000rcf for 30 seconds to separate suspended chitin and any bound cells from the medium. The supernatant was harvested, and the cell concentration was measured using qPCR.

### Measuring species abundance in chitin cocultures

Cultures were grown on colloidal chitin until early stationary phase, when the OD_600_ does not rise for at least two hours and all visible chitin has been consumed. Total CFU/mL are counted by plating the cultures onto MB2216 agar plates. The identity of 25 colonies derived from each culture were determined using qPCR, which is used to calculate the fraction of each species in the total CFU/mL.

### Exochitinase assay on particle surfaces

To compare exochitinase activity between the particle-bound and planktonic fraction, exochitinase activity was measured using an in *vitro* enzyme assay that relies on the enzymatic hydrolysis of 4-nitrophenyl N-acetyl-β-d glucosaminide to p-nitrophenol. *Vib*1A01 and *Psy*6C06 were grown on colloidal chitin for 24 hours and 500 μL of sample was centrifuged at 1000 rcf for 30 seconds. The supernatant was removed as the planktonic fraction. Fresh MBL was added back to the pellet, containing all bound cells and chitinases, to bring the volume back to 500 μL. A mastermix solution was prepared that contains 100mM sodium acetate and 2mM 4-nitrophenyl N-acetyl-β-d glucosaminide. 150 μL of this mastermix was added to 150 μL of the bound or planktonic fraction. The reaction was allowed to proceed for 30 minutes at room temperature. To quench the reaction, 100 μL of the enzyme reaction was centrifuged at max speed and the supernatant was added to 100 μL of 10% ammonium hydroxide, 2mM EDTA. The assay provides a colorimetric readout at 405nm. Pure p-nitrophenol is used as a standard to generate a calibration curve.

### Untargeted mass spectrometry

Metabolomics was performed using Liquid Chromatography coupled to an Agilent 6520 Time of Flight Quadrupole Time of Flight Mass Spectrometer in positive mode, 4GHz, high resolution mode. The column was an Agilent EC-CN Poroshell column (50×2.1mm, 2.7 μM), operated isocratically, which has been shown to reduce the interference of salts on metabolite ionization^61^. The buffer contains 10% Acetonitrile (Chromasolv), 90% water, and 0.01% formic acid. The flow rate was 350 μL/min. The sample was diluted 20 fold in MillQ water prior to measurement, and 3 μL was injected every 2 minutes. Raw data for all measurements was subjected to a spectral processing and alignment pipeline using Matlab (The Mathworks, Natick) as described previously^62^. Ions were annotated with a tolerance of 5mDa against a curated compound library that contains metabolites predicted to be in at least one species used in this study based on the BioCyc database^63^. Annotated metabolites and their intensities can be found in **supplemental dataset 1**. All raw spectral files have been deposited to the MassIVE database (MassIVE MSV000093541) with password reviewer123, and will be made publicly available upon manuscript acceptance.

### Fed batch reactors

Six parallel nutrient-limited fed batch reactors were set up as follows in a 25°C controlled room. First, 20ml of MBL medium with no carbon source was placed into a 250ml Erlenmeyer flask (bioreactor) containing a magnetic stir bar and a silicone sponge closure to minimize contamination (Sigma, C1046). Feed medium was placed in a separate bottle containing a pierceable rubber cap (Fisher, 15896921). Here, the feed medium contains MBL supplemented with 1mM GlcNAc. A feed tube was configured to transport medium between the feed bottle and the Erlenmeyer flasks, powered by a peristaltic pump (Ismatec IPC8). First, a stainless steel needle (18 gauge, 6inch stainless steel 304 syringe needle Sigma Z102717) withdraws medium from the feed bottle. This is connected to a PharMed BPT tubing (2.79mm) through a male leur fitting for 1/8in tubing (Sigma 21016). Next, this tube is connected to a PharMed 2- stop tubing 0.25mm (Ismatec 95723-12) using another 1/8in male leur fitting and a 22 gauge, 51mm metal hub needle (Hamilton HAM191022) that is inserted into the two-stop tubing. This two-stop tubing is placed through the peristaltic pump, where it is then connected to an additional length of 2.79mm PharMed BPT tubing and another 18 gauge stainless steel needle. This needle is then placed directly into the bioreactor, with the tip dispensing medium directly into the culture at a continuous flow of 0.3mL/hr. All species were inoculated into the bioreactors at an initial cell concentration of 1×10^7^ cells/mL from an overnight preculture. The preculture was first centrifuged and resuspended in MBL with no carbon to minimize any nutritional carryover.

### Coculture device

The membrane separated coculture device was manufactured according to the previously published design, CAD files, and assembly instructions^49^. This previous publication contains detailed instructions and video of how to assemble the device. The polycarbonate membranes (Isopore™ Membrane Filter, 0.1 μm VCTP; EDM Millipore) separated two cultures each containing 2.5ml of chitin MBL. All cells were inoculated at an initial density of 1×10^7^ cells/mL and sealed with an adhesive gas-permeable seal (Thermo AB-0718). The entire device was placed on an orbital mini shaker at 300RPM (VWR 444-0269) in a 25°C controlled room. OD_600_ was measured by withdrawing 100 μl from each culture, centrifuging at 1000rcf for 30 seconds, and measuring the OD_600_ of the supernatant in a 384 well plate.

### Protein functional prediction

Sequences of all proteins were obtained for each strain from NCBI (Table S1) and used as input for genome-wide functional prediction using InterproScan 5 REST API^64^. After functional prediction using default settings, results were merged and Pfam annotations were used to find any predicted domain annotations associated with beta-lactamases (PF00144, PF00753, PF04273, PF12706, PF13354, PF13483 or PF19583) or penicillin amidases (PF01804).

### Measuring GlcNAc consumption

To quantify GlcNAc, we employed an Agilent 6545 LC-QTOF-MS in negative mode with high-sensitivity slicer, using a scan rate of 2 Hz, a mass range of 50–1700 m/z, and a fragmentor voltage of 110 V. The drying gas flow was set to 10 L/min, nebulizer pressure to 35 psig, skimmer voltage to 65 V, and gas temperature to 320°C. Separation was achieved using an Agilent HILIC-Z Poroshell column (100 mm × 2.1 mm, 1.8 μm particles). Mobile phase A consisted of LC-MS grade water with 10% acetonitrile and 0.3% ammonium hydroxide, while mobile phase B contained 90% acetonitrile. Samples were diluted 20-fold in 50% acetonitrile, and 4 μL was injected. The gradient started with 0% phase A, increased to 20% over 1 minute, held for 30 seconds, and then returned to 0% for equilibration until 5 minutes. Quantification was performed using Agilent Quantitative software, based on the intensity of peaks compared to standards with known GlcNAc concentrations, using the chloride adduct (256.058 m/z) for detection.

### Exponential growth model

To characterize bacterial growth kinetics to estimate time to reach OD_600_ 0.25, we fit an exponential growth model to experimentally determined OD_600_ data for cultures in log phase. Log phase is the period of growth in which the log transformed growth data is linear for at least four consecutive time points with a goodness of fit (R^2^) greater than 0.95. The growth model is described by the equation:

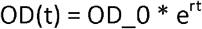

Where OD(t) is the OD600 at time t, OD_0 is the OD600 at the beginning of log phase growth, and r is the growth rate.

### Microscope images

Samples (200□μl) were taken from exponentially growing *Vib*1A01 or *Psy*6C06 colloidal chitin cultures at similar planktonic ODs (∼0.4). Chitin flakes were stained with 1□μg/ml FITC-WGA and cells were stained with a membrane dye FM 4-64 with a concentration of 5□μg/ml. The sample was then transferred to a glass bottom chamber for imaging with a Leica TCS SP8 inverted confocal microscope. The WGA signal was read in the GFP channel, which was excited with a 488□nm diode laser and the FM 4-64 was read in the mCherry channel and excited with a 580□nm diode laser. Fluorescence for both channels was detected through a ×40/1.3 objective and a highly sensitive HyD SP GaAsP detector.

### Chitinase purification and supplementation

*Psy*6C06 was streaked from -80°C glycerol stocks onto 1.5% agar plates containing MB 2216 (Fisher) medium. An overnight preculture was inoculated from a single colony in MB 2216. The following day, a 1% inoculum of this preculture was added to 400 mL of MBL medium with 2g/L colloidal chitin and stirred at 200 rpm until the culture reached the early stationary phase. Following growth, the culture medium was centrifuged at 2800 rcf for 20 minutes to remove residual chitin and cells, and then sterile filtered using a 0.2 μm membrane. The supernatant was concentrated 10-fold using an Amicon stirred cell (Millipore) with a 3 kDa cutoff filter. A protease inhibitor (Roche cOmplete EDTA-free protease inhibitor cocktail) was added to the final concentrate. The resulting solution was aliquoted into 500 μL portions in 1.5 mL microcentrifuge tubes, snap frozen in liquid nitrogen, and stored at -80°C until use. The protein concentration in the chitinase enzyme cocktail was determined to be 0.07 mg/mL using the Bradford reagent. When used to supplement cultures during growth on chitin, enzyme was added to the inoculum at a 1% concentration from this stock.

## Acknowledgements

We would like to sincerely thank Andreas Sichert and Otto X. Cordero for their valuable discussions, which significantly contributed to the advancement of this work.

## Supplemental information

**Table S1:** Species used in this study and NCBI accession numbers for genomic sequences.

| Species | Taxonomic ID | Functional Guild | NCBI BioProject | NCBI BioSample |
| --- | --- | --- | --- | --- |
| AltA3R04 | Alteromonas sp. | Exploiter | PRJNA478695 | SAMN19350919 |
| AmphC1R06 | Amphritea sp. | Scavenger | PRJNA478695 | SAMN19350932 |
| CitC3M06 | Citricella sp. | Scavenger | PRJNA478695 | SAMN19350936 |
| ColC2M11 | Colwellia psychrerythraea | Exploiter | PRJNA478695 | SAMN19350933 |
| MarD2M19 | Marinobacter sp. | Scavenger | PRJNA478695 | SAMN19350941 |
| MarF3R08 | Marinobacter sp. | Scavenger | PRJNA478695 | SAMN19350954 |
| MarF3R11 | Marinobacter sp. | Scavenger | PRJNA478695 | SAMN09522136 |
| ParaC2R09 | Paracoccus kamogawaensis | Scavenger | PRJNA478695 | SAMN19350935 |
| PhaB3M02 | Phaeobacter sp. | Exploiter | PRJNA478695 | SAMN09522130 |
| PseuI2R16 | Pseudoalteromonadaceae | Exploiter | PRJNA478695 | SAMN19350960 |
| Psy6C06 | Psychromonas sp. | Degrader | PRJNA414740 | SAMN08130274 |
| ReinG2M07 | Reinekea sp. | Scavenger | PRJNA478695 | SAMN19350955 |
| SilA3R06 | Silicibacter sp. | Exploiter | PRJNA478695 | SAMN19350920 |
| Vib1A01 | Vibrio splendidus | Degrader | PRJNA414740 | SAMN07809270 |
| Vib6D03 | Vibrio penaeicida | Degrader | PRJNA414740 | SAMN08130373 |
| VibC3R12 | Vibrio sp. | Exploiter | PRJNA478695 | SAMN09522132 |
| VibG2R10 | Vibrio sp. | Degrader | PRJNA478695 | SAMN19350956 |
| VibI3M07 | Vibrio sp. | Degrader | PRJNA478695 | SAMN19350961 |

**Table S2:** CFU/mL/OD values for each species grown in MB2216 precultures. These values are used to determine the dilution required for precultures before inoculating experimental cultures that require consistent cell densities such as in Fig. 1A and 3E.

| **Species** | **CFU/ml/OD** |
| --- | --- |
| **AltA3R04** | 1.71E+09 |
| **AmphC1R06** | 2.72E+08 |
| **ColC2M11** | 7.01E+08 |
| **MarD2M19** | 4.04E+08 |
| **MarF3R08** | 1.14E+09 |
| **MarF3R11** | 1.30E+09 |
| **ParaC2R09** | 7.63E+08 |
| **PhaB3M02** | 7.23E+08 |
| **Psy6C06** | 5.06E+08 |
| **SilA3R06** | 1.89E+09 |
| **Vib1A01** | 4.15E+08 |
| **VibC3R12** | 3.56E+08 |
| **VibG2R10** | 3.52E+08 |
| **VibI3M07** | 8.32E+08 |

**Table S3:** Lag times (in days) of mono- and cocultures of degraders with exploiters (top) or scavengers (bottom). NA denotes cultures that showed no growth. Red boxes are cocultures with a delayed lag time compared to the monoculture. Green boxes are those with a decreased lag time compared to the monoculture.

| Degrader | Monoculture | Exploiter |  |  |  |  |
| --- | --- | --- | --- | --- | --- | --- |
|  |  | AltA3R04 | ColC2M11 | PhaB3M02 | SilA3R06 | VibC3R12 |
| Vib1A01 | 1.9 | 1.9 | NA | 1.9 | 1.9 | 3.51 |
| VibG3R10 | 2.88 | 3.83 | NA | 4.9 | 2.88 | 4.68 |
| VibI3M07 | 1.9 | 2.88 | NA | 2.88 | 3.83 | 4.9 |
| Psy6C06 | 4.1 | 4.72 | 4.36 | 4.14 | 4.94 | 4 |

| Degrader | Monoculture | Scavenger |  |  |  |  |
| --- | --- | --- | --- | --- | --- | --- |
|  |  | AmphC1R06 | MarD2M19 | MarF3R08 | MarF3R11 | ParaC2R09 |
| Vib1A01 | 1.33 | 1.33 | 2 | 2 | 2 | 1.1 |
| VibG3R10 | 2.65 | 3.13 | 5.46 | 3.13 | 2.81 | 4.71 |
| VibI3M07 | 2.65 | 2.19 | 3.13 | 2.5 | 3.13 | 3.13 |
| Psy6C06 | 4.21 | 4 | 4.42 | 4.14 | 5.33 | 4.42 |

**Table S2:**
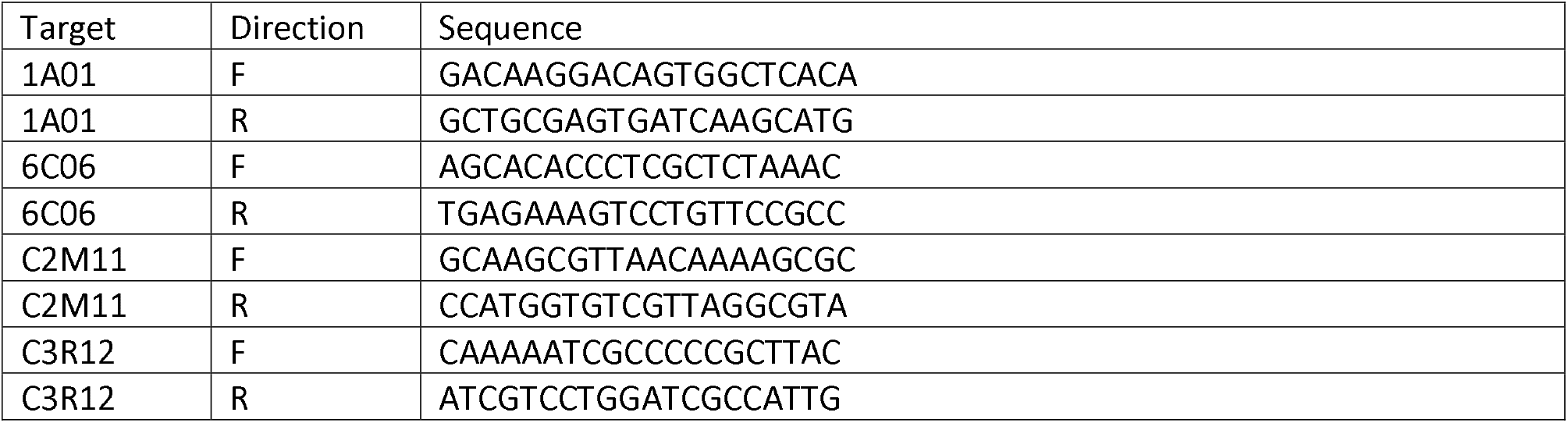
qPCR primers used in this study

**Figure S1:**
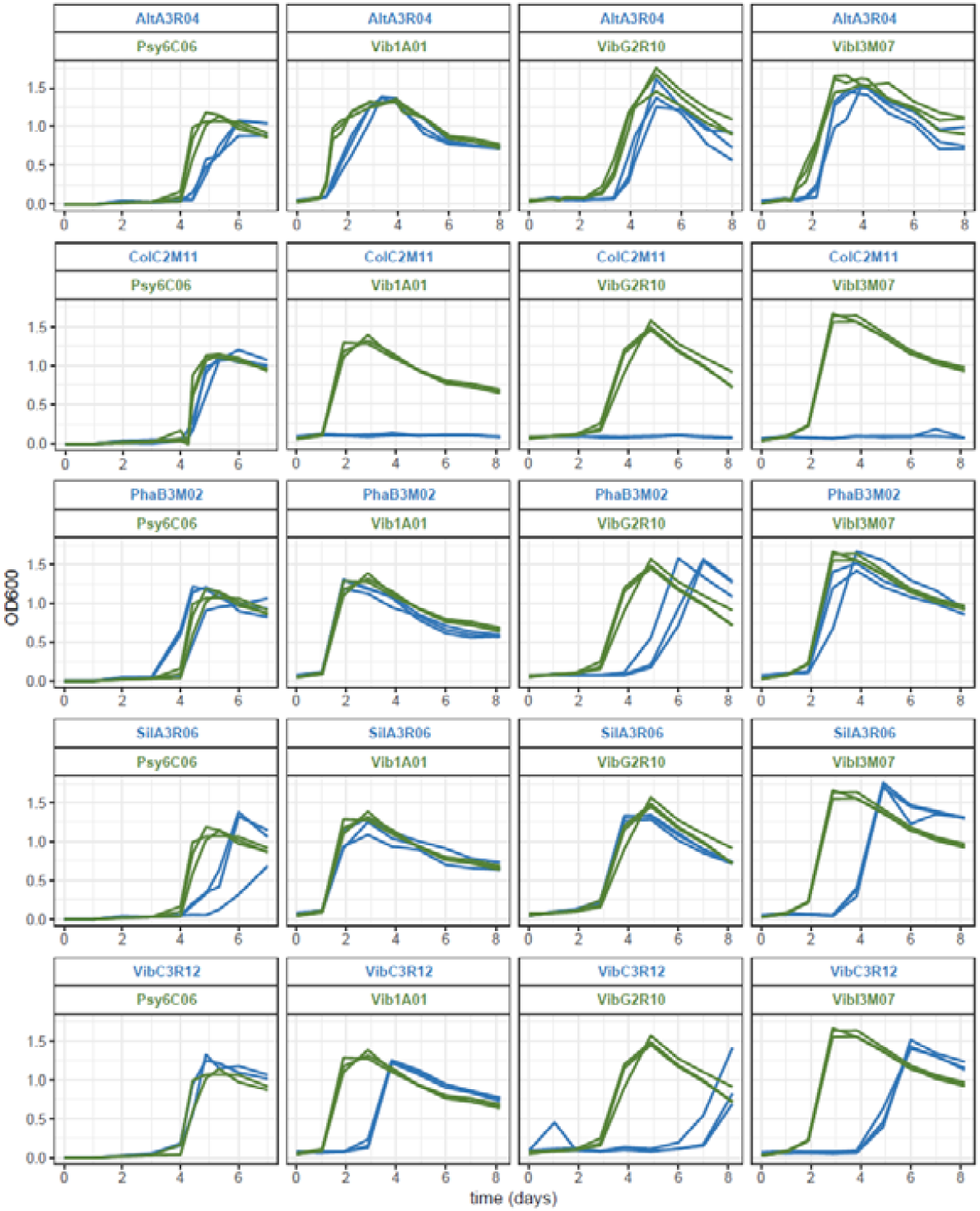
Growth of pairwise cocultures of each degrader (green text), and exploiter (blue text) on chitin as a sole carbon source. Green lines represent growth of degrader monocultures, blue is degrader:exploiter cocultures. All three biological replicates are shown. Each species is inoculated at a density of 10^7^ cells/mL from a preculture of MB2216. The exact volumes needed are calculated using OD_600_ to cell density values from Table S2.

**Figure S2:**
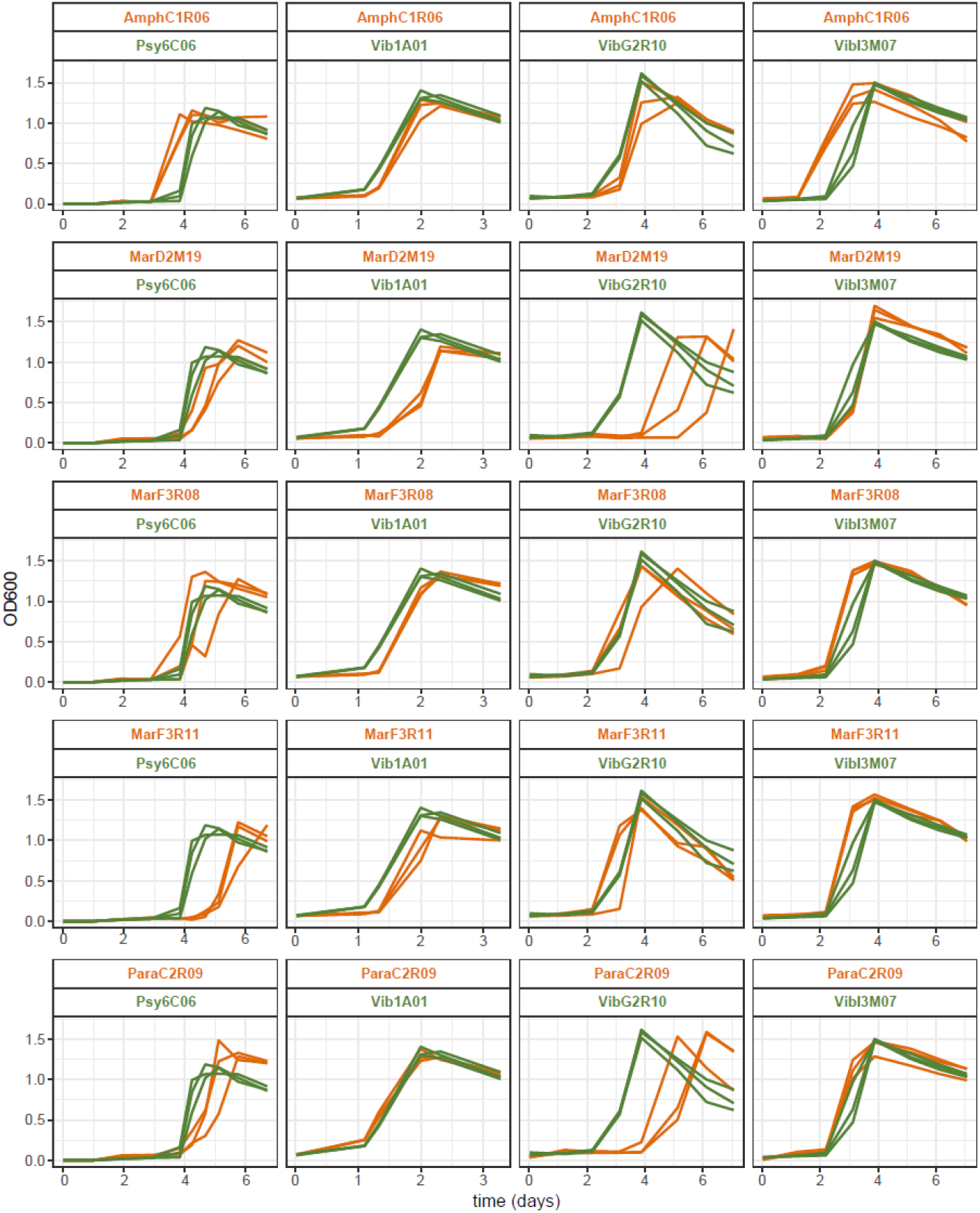
Growth of pairwise cocultures of degraders (green text), and scavengers (orange text) on chitin as a sole carbon source. Green lines represent growth of degrader monocultures, orange is degrader:scavenger cocultures. All three biological replicates are shown. Each species is inoculated at a density of 10^7^ cells/mL from a preculture of MB2216. The exact volumes needed are calculated using OD_600_ to cell density values from Table S2.

**Figure S3:**
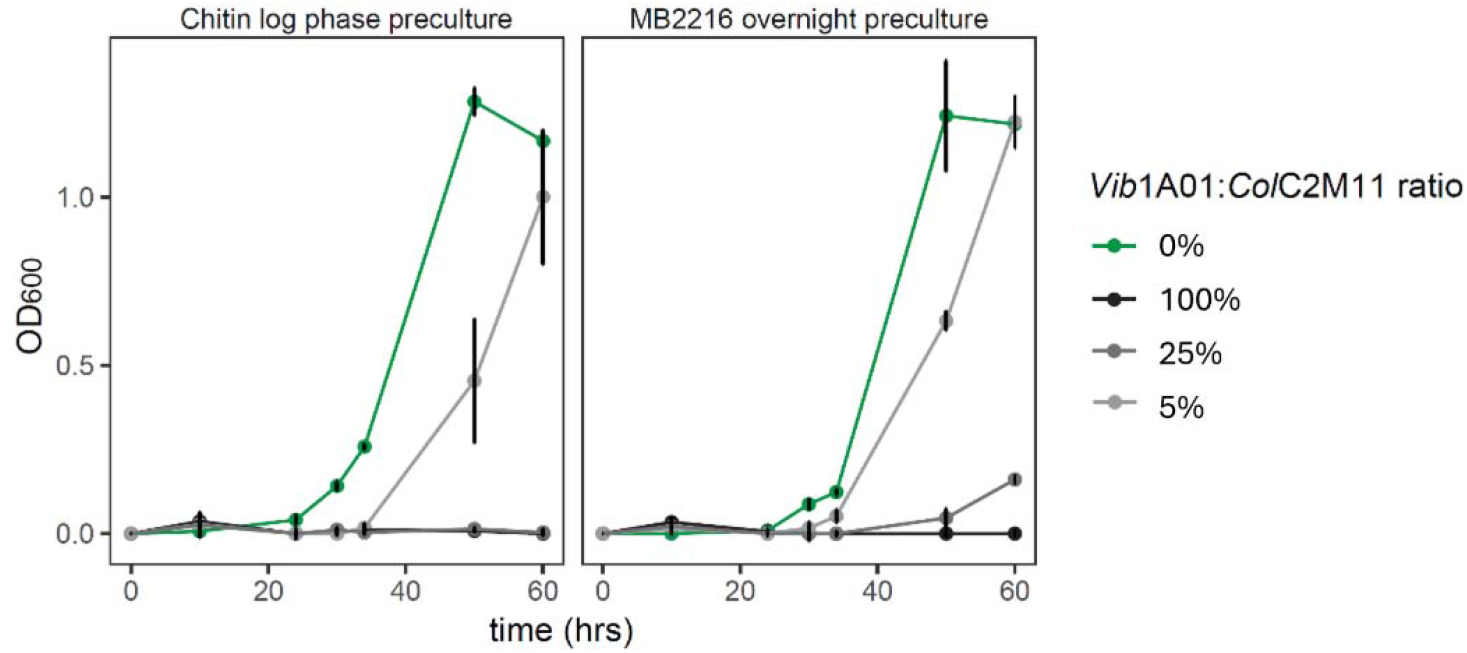
Growth on colloidal chitin of *Vib*1A01 monoculture or coculture with *Col*C2M11 with different inoculation ratios or preculture conditions. Precultures were taken from *Vib*1A01 growing in MB2216 rich medium overnight culture or from log phase growth on colloidal chitin. In all cases, *Col*C2M11 is taken from a MB2216 overnight preculture. Experiments were performed in triplicate, and error bars represent standard deviation from the mean. In all cases, *Vib*1A01 is inoculated at 1×10^7^ cells/mL.

**Figure S4.**
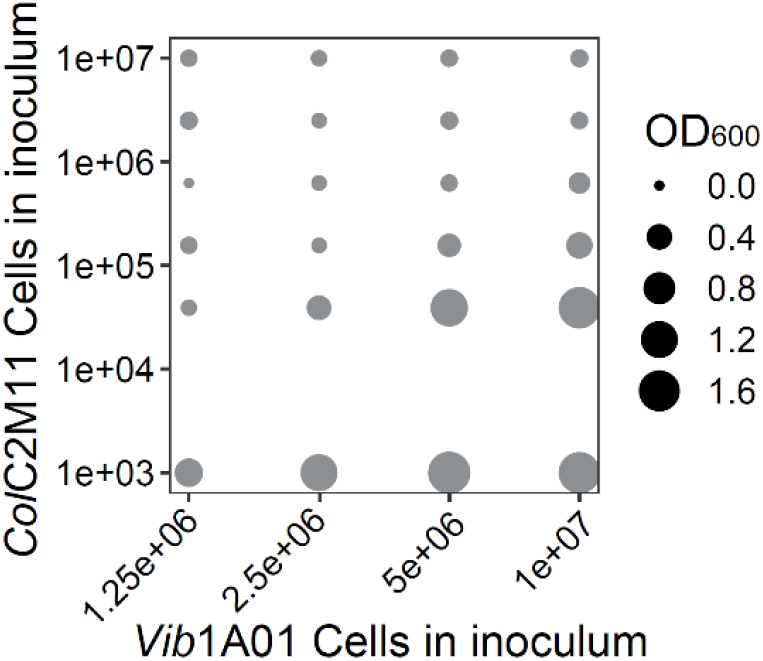
Growth of *Vib*1A01 cocultures with *Col*C2M11 on colloidal chitin at different initial cell ratios at 44 hours. The experiment was performed in triplicate.

**Figure S5:**
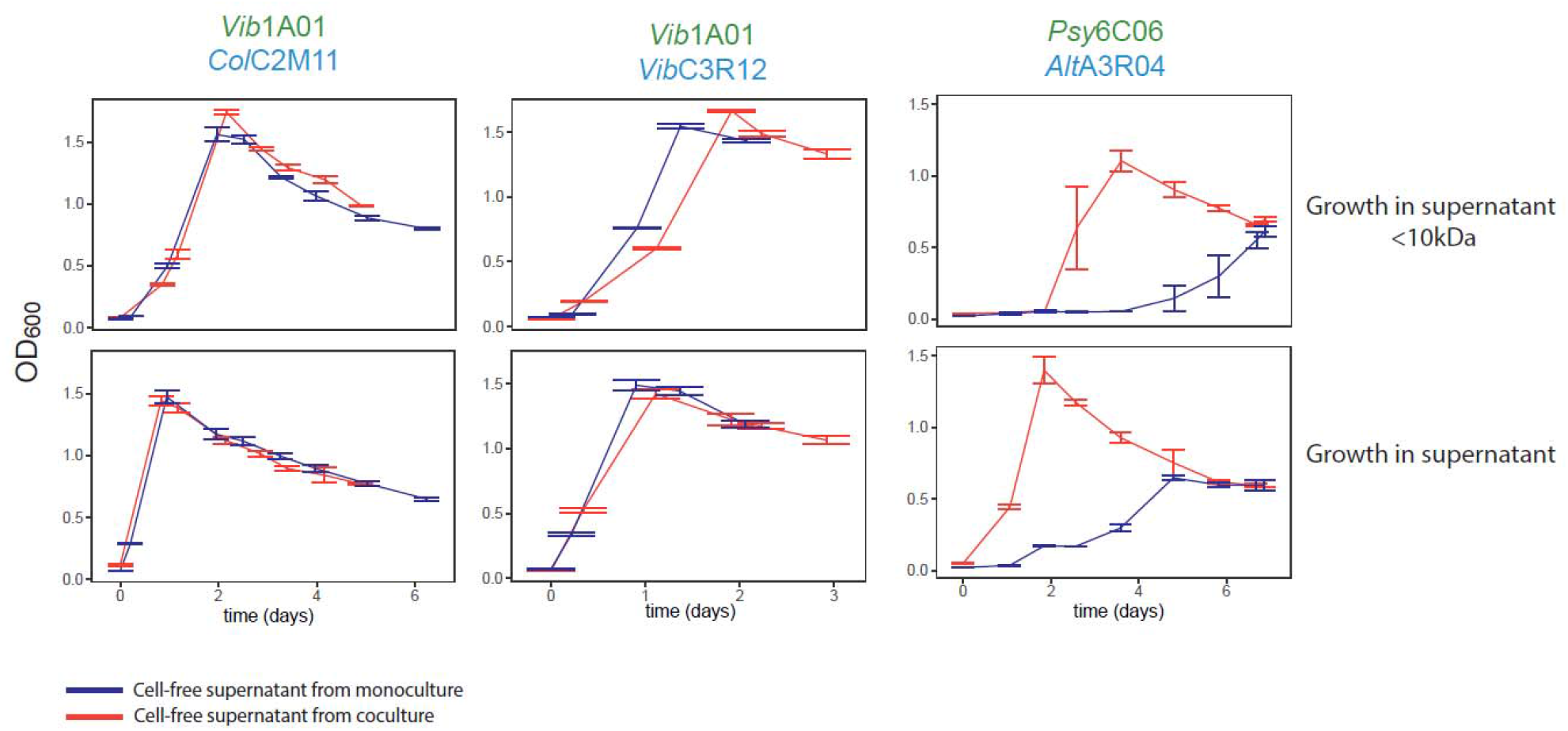
Growth of degraders (green text) in cell-free supernatants or in supernatants with large compounds (10kDA) removed. Supernatants originate from cocultures of the degrader in coculture with the respective exploiter (blue text). Error bars represent standard deviation of the mean of triplicates.

**Figure S6:**
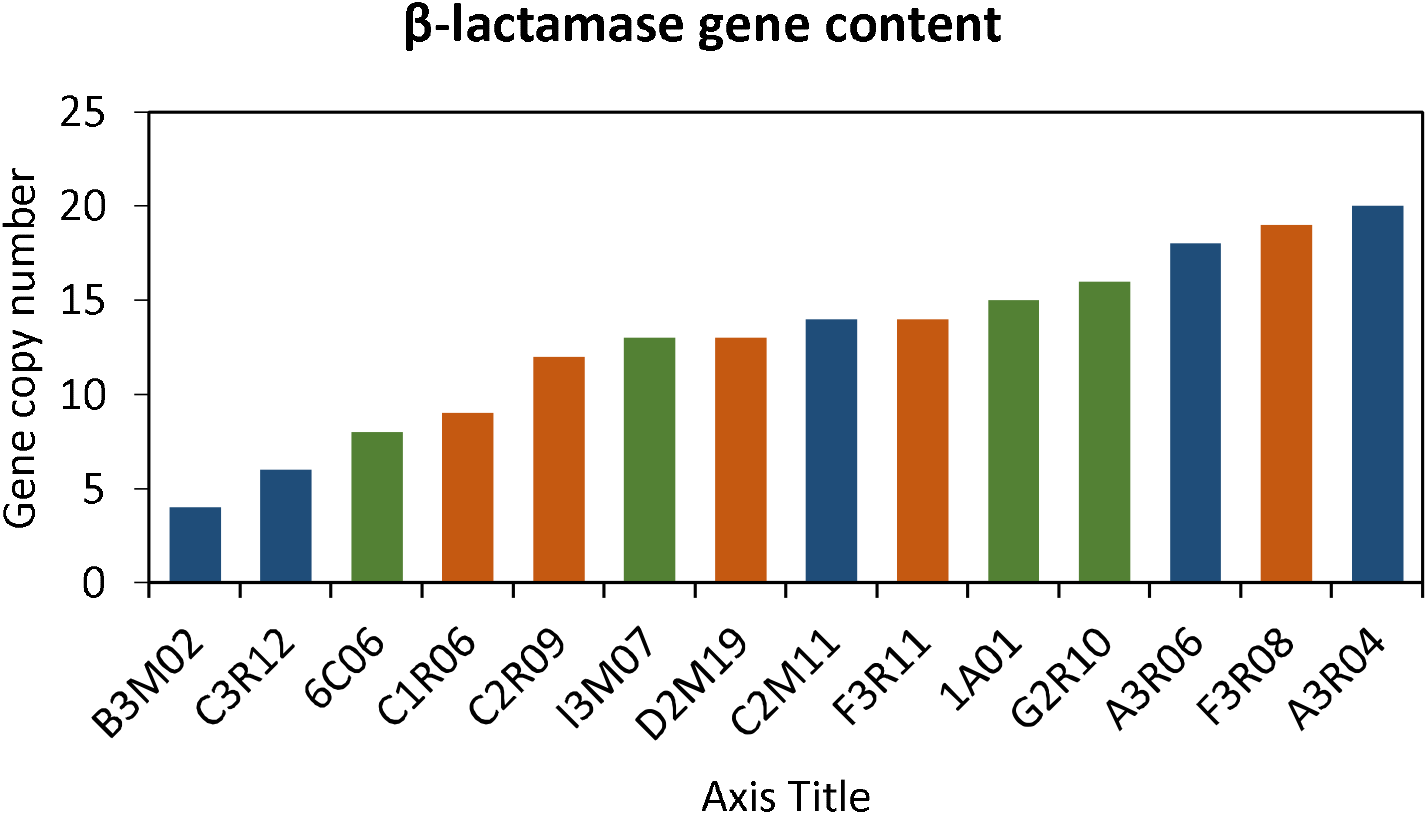
Copy number of β-lactamases per genome in each species used in this study. Degraders are in green, exploiters are in blue, scavengers are in orange.

### Supplementary Note 1: Nutrient competition in fed-batch culture

In this Note, we analyze the competition of two species growing on the same nutrient, GlcNAc, in a fed-batch culture. The result will be used to interpret the data of the fed-batch co-culture of *Vib*1A01 and *Col*C2M11 as described in the main text.

#### 1. Solution for the monoculture

We first describe the monoculture in the fed-batch reactor.. Let *ρ* (*t*) be the biomass density (or OD) of a single species of bacteria grown in the fed-batch culture, inoculated at *ρ*_0_at time *t* = 0. Let the nutrient concentration in the culture be *n* (*t*), and the nutrient drip rate be *α*. Then the dynamics of the system are described by the following equations:

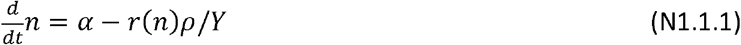

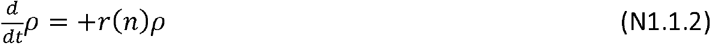

where *Y* is the biomass yield of the nutrient, and

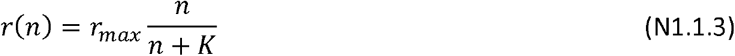

the Monod growth kinetics, with saturated growth rate *r*_*max*_, and Monod constant *K*.

Eq. (N1.1.1) and (N1.1.2) give the condition,

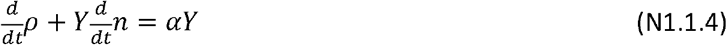

reflecting the constraint that the added nutrient amount is either converted to biomass or remains in the culture. If most of the nutrient is captured by the cells (as will be shown shortly below), then 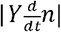 is negligible compared to 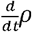, and Eq. (N1.1.4) is easily solved, giving

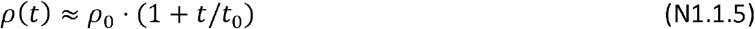

Where *t*_0_ ≡ *ρ*_*0*_ /(*αY)*. Inserting the approximate solution (N1.1.5) into Eq. (N1.1.2) gives the dynamics of *n* (*t*) :

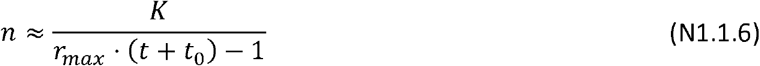

Since 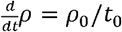 and 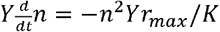, then the approximation 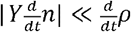 requires that

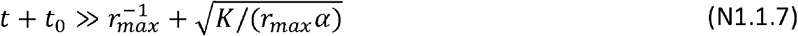

For *rmax* ∼ 0.5/*h,K* ∼ 1 *μM*, and with the drip rate being *α* ∼10*μM* /*h*, the right hand side of the inequality is ∼1 *h*. For an initial inoculant at *ρ*_*0*_ OD and an (inverse) yield being *Y*^−1^ = 6 mM GlcNAc/OD, we have *t*_0_∼ 6 *h*. Thus, the condition (N1.1.7) is satisfied for all, and the solutions (N1.1.5) and (N1.1.6) are almost exact.

From Eq. (N1.1.6), we note also that 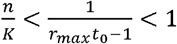. Hence, the Monod growth kinetics (N1.1.3) can be approximated by its linear form, i.e., *r(n)* ≈ *r*_*max*_ *n/K*. This is the form we will use below. To simplify the notation, we define;*V* ≡ *r*_*max*_ */K* such that *r(n)* ≈*v* · *n*.

#### 2. Two-species competition

Now we consider two species (A and C), with biomass *M*_*A*_ *(t)* and *M*_*C*_ *(t)*, respectively, whose growth rates are (in the linear approximation) *r*_*A*_*(n)* ≈*v*_*A*_ · *n, r*_*C*_ *(n)* ≈*v*_*C*_ · *n* with *V*_*A*_≡ *rmax /K*_*A*_, and *V*_*c*_≡ *rmax /K*_*C*_. The equations describing the two species dynamics are

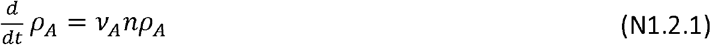

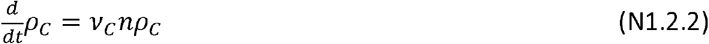

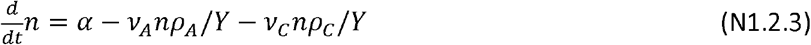

where the same yield *Y* is taken for the two species for simplicity.

The total biomass density *ρ ≡ ρ*_*A*_*(t)* + *ρ*_*C*_*(t)* again satisfies the condition (N1.1.4), and with 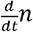 negligible, its solution is again the linear form given in Eq. (N1.1.5). To see how biomass is partitioned between the two species, we define the fractional biomass, *ϕ*_*i*_ *≡ ρ*_*i*_*(t) ρ(t)*. In terms of *ϕ*_*i*_ the dynamics of *ρ* can be written (from Eqs. (N1.2.1) and (N1.2.2)) as

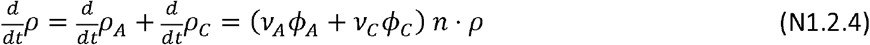

and the dynamics of *ϕ*_*i*_ are given as

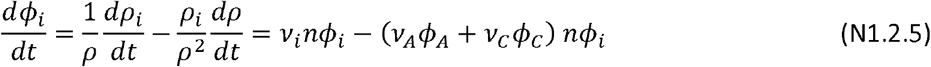

where we used Eqs. (N1.2.1), (N1.2.2) and (N1.2.4) in the last step above. Using *ϕ*_*A*_ +*ϕ*_*C*_ *=*1, Eq. (N1.2.5) can be written as

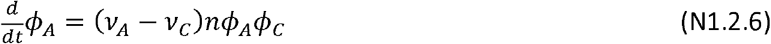

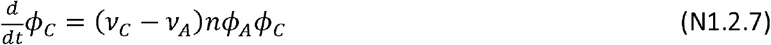

or in terms of the fractional difference, Δ*ϕ* ≡ *ϕ*_*C*_ *− ϕ*_*A*_

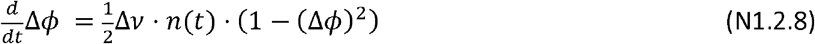

Where Δ*V* ≡ *V*_*C*_ *− V*_*A*_.

To solve Eq. (N1.2.8) and obtain the dynamics of Δ*ϕ*, we need to know the form of *n*(*t*). This can be obtained by exploiting Eq. (N1.2.4). Writing 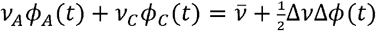 where 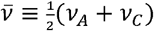, we have

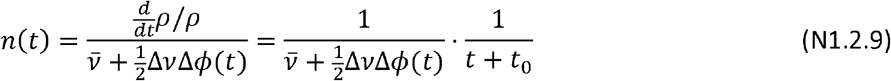

where we used the solution (N1.1.5) for *ρ(t)*. Inserting this form of *n*(*t*) in Eq. (N1.2.8), we obtain a closed ODE for Δ*ϕ* (*t*),

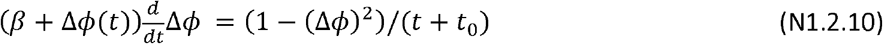

With 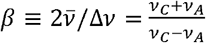.

Eq. (N1.2.10) can be integrated exactly:

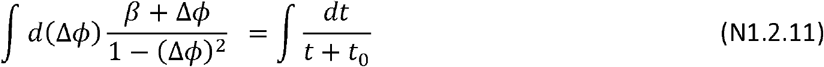

leading to the solution (for initial condition Δ*ϕ(0) =0*, i.e., *ϕ*_*A*_ *(0) =ϕ*_*C*_ *(0) = 0.5)*:

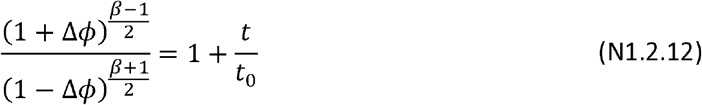

or expressed more transparently as

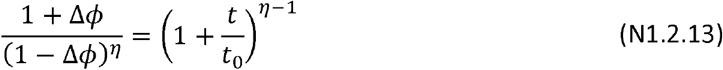

Where *η ≡ V*_*C*_ */ V*_*A*_, a dimensionless ratio of the Monod parameters, gives the ratio of growth rates (i.e., *η = r*_*C*_ */ r*_*A*_) at low nutrient concentrations where *r*_*i*_*= r*_*i*_ *n*.

To see the form of the solution described by Eq. (Nsss1.2.13), let us suppose *η*>1. Then in the long time limit (*t ≫ t*_0_) where the right-hand side diverges, the left-hand side must also diverge, which happens for Δ *ϕ* =*ϕ*_*C*_ −*ϕ*_*A*_ → 1 (corresponding to *ϕ*_*C*_ → 1and *ϕ*_*A*_ → 0). The asymptotic solution can be written as Δ *ϕ (t)* ≈ 1 − 2 ^1/ *η*^ (*t*/*t*_0_) ^−(*η* −1)/ *η*^, or 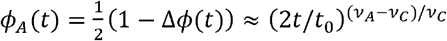 i.e., the slower-growing species, in this case, species A because we took *η>* 1 (*V*_*C*_> *V*_*C*_), is depleted from the co-culture in a power-law fashion, with an exponent that is proportional to the difference in the growth parameters.

Alternatively, the limit that *η* is slight larger than 1, i.e., for *η* −1≪1, such that Δ *ϕ* ≪1 also, Eq. (N1.2.13) can be simplified to

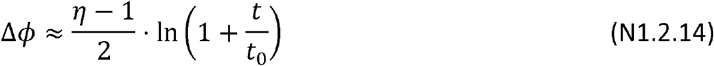

#### 3. Quantitative comparison to experimental results

The calculation in the previous sections assumed that the volume of the culture is unchanged. However, to compare to the experimental data quantitatively, we need to incorporate the fact that in the fed-batch reactor, the addition of GlcNAc does change the culture volume. According to **Method**, the flow to the bioreactor is *α*_*V*_ = 0.3*ml/h*. For a culture with initial volume of *V*_0_= 20 *ml/h*, the volume changes as *V(t) = V*_*0*_+*α*_*V*_ (*t*), with an increase of ∼20ml over the course of the 72 h experiment. The volume change affects the GlcNAc concentration *n* (*t*) and cell densities *ρ*_*i*_ (*t*). To treat this effect, we introduce the GLcNAc amount *m* (*t*), with *n* (*t*) = *m* (*t*)/*V (t)*, and the biomass of each species, *M*_*i*_ (*t*), with *ρ*_*i*_ (*t*) = *M*_*i*_ (*t)*/*V*(*t)*. Eqs. (N1.2.1)-(N1.2.3) are then modified to

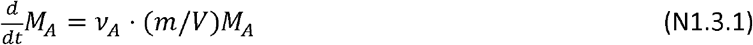

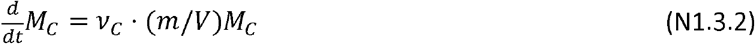

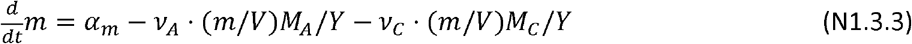

Since GlcNAc inflow is at 1mM, the rate of GlcNAc increase is *α*_*m*_= 1*mM*. 0.3*mLh*^−1^ = 0.3 *μmol h*^−1^ The mass conversation condition becomes 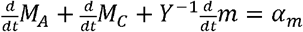. Neglecting 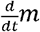 as before, we obtain a simple relation for the total biomass *M* (*t*) ≡ *M*_*A*_ (*t*) + *M*_*C*_ (*t*)

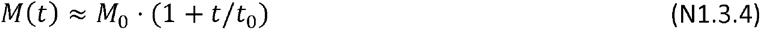

Where *t*_0_ ≡ *M*_*0*_*/(α*_*m*_ *Y*). For the coculture with initial inoculant of each species at OD = 0.01 and an initial reaction volume *V*_0_ = 20 *ml*, we have *M*_*0*_ = 0.4 OD×ml. This gives *t*_*0*_ = 8*h*.

The dynamics of *ϕ*_*i*_ = *M*_*i*_/ *M* can be derived as in Sec. 2, leading to Eqs. (N1.2.6)-(N1.2.8), with *n* (*t*) replaced by *M* (*t*)/ *V* (*t*), and with

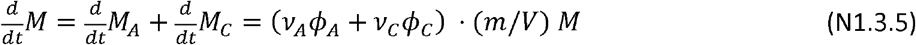

Then, Eq. (N1.3.5) can be used to solve for the factor *m*/*V*, with

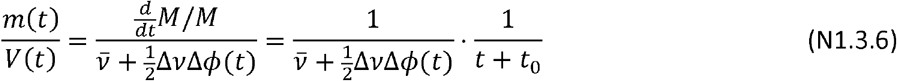

Thus, the final equation for Δ *ϕ* is the same as Eq. (N1.2.10), with the same solution (N1.2.13) [and its approximation (N1.2.14)], except that the value of *t*_0_ is given by *ρ*_0_ *V*_0_/*(α*_*m*_ *Y*); in other words, the GlcNAc influx rate *α* of the constant volume scenario considered in Sec. 1 and 2 is replaced by *α*_*m*_ */V*_0_.

In **Fig. N1a**, we plot the dependence of Δ *ϕ*vs *η* according to solution Eq. (N1.2.13) for several values of the fed-batch runtime *t* and for *t*_0_ = 8*h* (solid lines). For *t* = 72*h* (black line), we compare the analytical solution to the exact numerical solution calculated from Eqs. (N1.3.1)-(N1.3.3) (black circles). The analytical solution is seen to be in quite good agreement with the exact numerical solution. **Fig. N1b** shows a zoomed in view for nutrient uptake ratio *η* ≈ 1. The dashed line is the approximate solution given in Eq. (N1.2.14).

**Figure N1:**
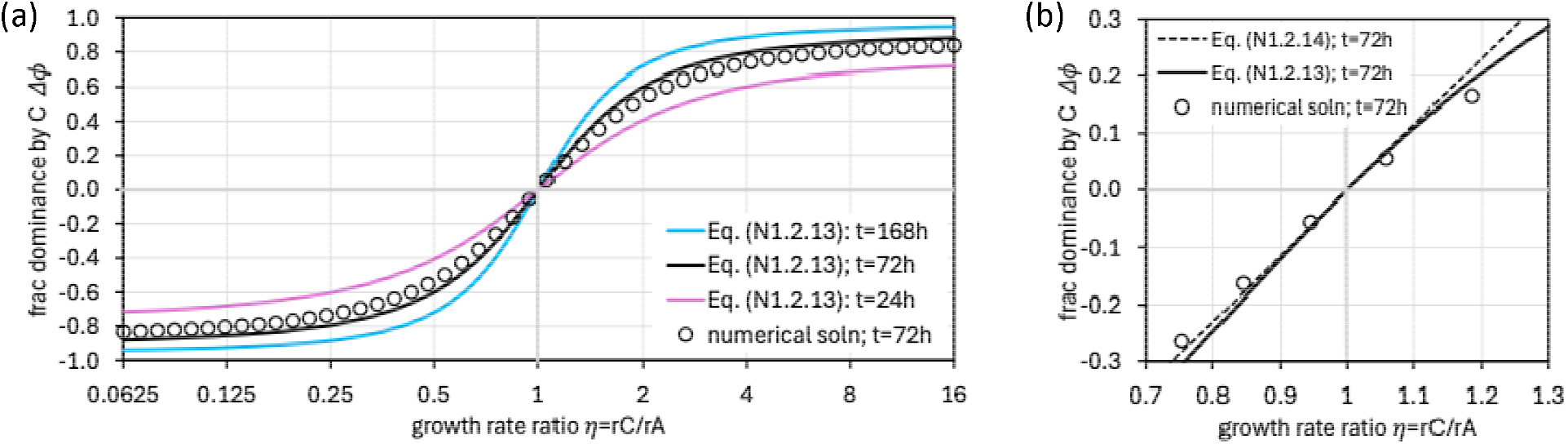
Comparing the analytical and numerical solutions of the system as defined by Eq. (N1.3.1)- (N1.3.3). Plotted is the dependence of the fractional species dominance Δ *ϕ ≡ (M*_*C*_ − *M*_*A*_) /*(M*_*C*_ + *M*_*A*_) for various values of η ≡ (*r*_*max,C*_/ *K*_*C*_)/ (*r*_*max,A*_/ *K*_*A*_, which gives the growth rate ratio *rC* :r*A* at low nutrient concentrations. **(a)** Solutions for various duration of fed-batch growth *t*, with *t*_0_ = 8*h* Solid lines of different colors show the analytical solution Eq. (N1.2.13), which can be rewritten as *η In[(1 +t/t*_*0*_) (1 + Δ *ϕ)/In[(1+t/t*_*0*_) (1 − Δ *ϕ)/*. Note that the solution exhibits the built-in symmetry that for, Δ *ϕ → −* Δ *ϕ, η* ≈ 1/ *η*, which arises from switching the species labels A and C. Open circles are the result of direct numerical solution. **(b)** Solution for systems with comparable uptake (*η ≈*1) after 72h of fed-batch growth, with the approximate solution Eq. (N2.1.14) shown as the dashed line.

Finally, we use the solution to extract the value of *η* from our fed-batch data: The data in **Fig. 3B** of the main text shows that the abundance of Vib1A01 is about 2/3 of that of ColC2M11 at 72h, i.e., 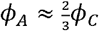; this leads to *ϕ*_*A*_ ≈40%,*ϕ*_*C*_ ≈ 60%, or Δ*ϕ*, ≈ 20%. For *t* = 72 *h, t*_0_ =8 *h*, Eq. (N1.2.14) becomes. Then for ≈1, we have *η* ≈1; see **Fig. N1b**.

### Supplementary Note 2: Effect of nutrient competition on chitin degradation

In this Note, we describe a mathematical model describing the competitive metabolic interactions in a co-culture between a chitin degrader (*Vib*1A01, species A), which produces and excretes chitinases, resulting in the release of labile nutrients (GlcNAc), and an exploiter (*Col*C2M11, species C), which competes with the degrader in the uptake of the labile nutrient. Our goal is to demonstrate that a metabolism-based model can recapitulate the drastic effect observed in the main text: that the addition of a small amount (10%) of the exploiters can inhibit growth (**Fig. 1D** of the main text) even though the exploiters do not exhibit much stronger affinity for GlcNAc (Supp. Note 1 and Figure 3). We will show that the sensitivity to exploiters is possible in the context of an incipient Allee transition for the monoculture. In this note, we follow the model and notation introduced in Guessous et al.^34^ describing the coupled dynamics of the degraders, chitinases, and labile nutrients, but include the additional effect of nutrient uptake by the exploiters.

#### 1. Nutrient dynamics

We start by writing down the flux balance equation describing the generation, uptake, and diffusion of the labile nutrient (GlcNAc), whose concentration is denoted by *n* (*R,t*) around the surface of a spherical chitin particle of radius *R*_0_:

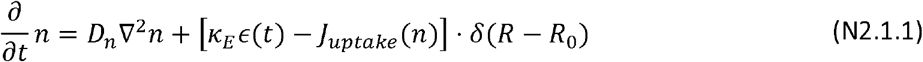

Here, *R* is the distance from the center of the chitin particle, *D*_*n*_ is the nutrient diffusion coefficient, *ϵ (t)* is the chitinase amount per surface area, and *K*_*E*_ is the rate of nutrient generation per chitinase (so that *K*_*E*_ *ϵ* is the nutrient generated by chitinases per surface area). The 2^nd^ term in the square bracket is the total flux of nutrient uptake by cells on the particle surface. Describing the replication rate of each species *i* by the Monod growth kinetics, *r*_*i*_(*n*) = *r*_*max,i*_ *· n/(n + K*_*i*_) where *r*_*max,i*_ is the batch culture growth rate and *K*_*i*_ is the Monod constant, then

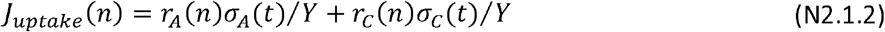

where *σ* _*i*_ (*t*) is the number of particle-attached cells of species *i* per surface area and *Y* is the biomass yield (taken to be the same for the two species).

Several key approximations are taken in the formulation of the dynamics:

- Nutrient uptake occurs only on the particle surface; this is justified by the drop in nutrient concentration away from the particle and is well-established numerically and experimentally in Guessous et al.^34^
- Cells and chitinases are assumed to be distributed uniformly across the particles (so that *σ* _*i*_ and *ϵ* are independent of the location on the particle). This assumption, taken for mathematical convenience, may be justifiable when both cell and chitinase densities increase. However, it is an oversimplification during the early stages when there are only a few cells on particles.

Eqs. (N2.1.1) and (N2.1.2) are supplemented by the no-flux boundary condition at the particle surface,

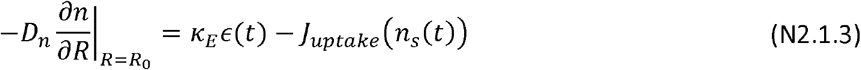

where *n*_*s*_(*t*) = *n(R*_*0*_, t) is the nutrient concentration at the particle surface, whose value determines the replication rates of the degraders and exploiters on the surface of the chitin particles, *r*_*A*_*(n*_*s*_) and r_c_(*n*_s_) respectively.

Since the dynamics of nutrient production and uptake happen on timescales, much faster than those of enzyme production and cell growth, we take the approximation that 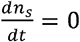 and look for the quasisteady-state solution *n*_*s*_ for given values *of σ* _*A*_, *σ* _*C*_, and ϵ: Since the replication rates *r*_*A*_, *r*_*c*_ are much below the batch culture growth rates *r*_*max,A*_, *r*_*max,C*_, respectively, the Monod kinetics can be approximated by the linear form*r*_*i*_(*n*) = *r*_*max,i*_ ·*n/K*_*i*_. In this case, the steady-state solution to Eq. (N2.1.1) becomes the 1/R-Coulomb potential, *n(R, t) = n*_*s*_*(t) · R*_*o*_*/R*, and the boundary condition Eq. (N2.1.3) leads to an important relation between *n*_*s*_ and the values of *σ* _*A*_, *σ* _*c*_, *ϵ:*

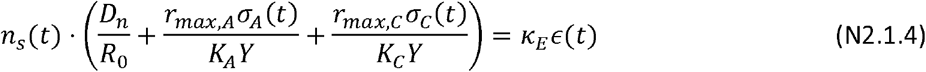

The time dependence indicated in these quantities refers to their inter-relationship as their quantities change over the scale of the cell doubling time. Note the distinction from the 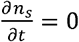 limit taken for Eq. (N2.1.1), which refers to the time scale of nutrient diffusion.

In Eq. (N2.1.4), the first time can be identified as the flux of nutrient loss, or leakage, due to diffusion, whose unit is the number of GlcNAc molecules per surface area per time. Note that this term is proportional to *D*_*n*_ as expected of diffusive loss, and inversely proportional to *R*_*0*_, reflecting a dependence on local geometry: the larger the curvature, the larger the leakage. To make contact with experimentally relevant quantities, it is helpful to convert the above formulation on a single chitin particle to bulk quantities involving the number of chitin particles (assumed to be of identical size *R*_*0*_) per volume, *ρchitin-* Let the chitinase “concentration” be 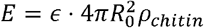, and the surface-attached cell density be 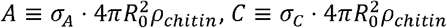. Multiplying 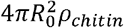 to the two sides of Eq. (N2.1.4), we obtain

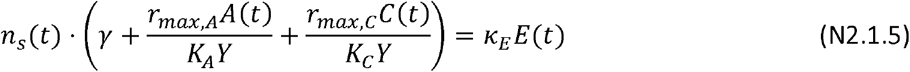

with a leakage rate given by 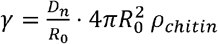. Let 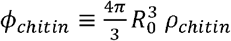.*be* the chitin volume fraction, given by the total chitin amount through the chitin density *d*_*chitin*_ ≈ *1.4lg/cm*^*3*^. For a total amount of 0.2%w/v, we find that ϕ_c*hitin*_ · *d* _*chitin*_ *= 0-002g/cm*^*3*^, or that the volume fraction ϕ _*chitin*_ ≈ 0.14%. For fixed *ϕ*_*chitin*_, we see that the leakage rate

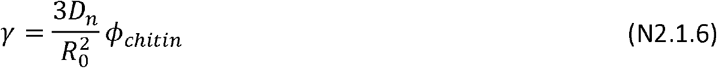

is inversely proportional to 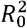

Comparing the situation studied by Guessous et al.^34^ (chitin chips where *R*_*0*_ ≈ 150 *μm)* to the current work (colloidal chitin particles where *R*_*o*_ ≈ 10 *μm)*, we find that the ratio of radii being 15x results in a leakage rate, *γ*, which is 225x larger for colloidal chitin for the same chitin density *ϕ* _*chitin*_ *-* Numerically, for 0.2%w/v chitin used in the current work,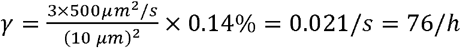. whereas for chitin chips in Guessous et al.^34^, *γ*, is only 0.3/h.

Eqs. (N2.1.5) and (N2.1.6) are the main take away of this analysis of nutrient flux balance at the particle surface. Eq. (N2.1.5) is a condition relating the nutrient concentration on the surface to the density of cells and enzymes, and Eq. (N2.1.6) gives the magnitude of the nutrient leakage ratey γ. The nutrient concentration *n*_*s*_*(t)* is fixed by macroscopic balances through the population growth rate as discussed below.

For a growing culture, *A*(t) and *E(t)* increase exponentially in steady state growth, and the nutrient leakage term *γ* is negligible. However, nutrient leakage can be important at low chitinase and cell densities during the initial phase of growth on chitin, especially for small chitin particles where the leakage rate *γ* is large. It is useful to re-express Eq. (N2.1.5) in a way that makes the leakage rate *γ* more directly meaningful. Factoring out a factor *r*_*max,A*_*/(K*_*A*_*Y)* from Eq. (N2.1.5), we have

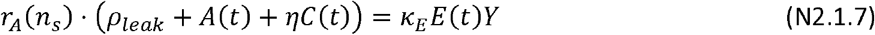

Where

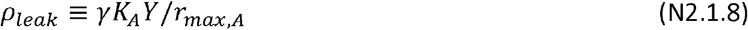

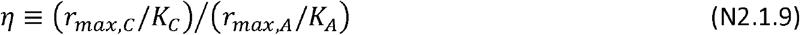

The term *ρ*_*leak*_ in Eq. (N2.1.7) expresses the leakage term as an effective cell density. It can be thought of as an additional population of degraders that the enzymes need to support through their generation of labile nutrient (even in the absence of the exploiters). It gives the scale of the exploiter density above which leakage can be neglected.

Numerically, taking the Monod constant^◊^ to be *K*_*A*_ ≈ 1 *μM,r*_*max,A*_ ≈ *0.6/h*., and the GlcNAc yield as 1 OD/ml/6mM **(Table N2)**, we obtain *p*_*leak*_ ≈ 0.02 *OD*, or 2 × 10^7^ cells/ml, for the colloidal chitin particles. Thus, the leakage flux is a substantial factor that needs to be considered for the range of cell densities that we study (where the initial inoculant used was 0.01 OD). Finally. Eq. (N2.1.7) also shows that the presence of the exploiters, at density *C*, serves effectively as an additional leakage term that increases *ρ*_*leak*_ by an amount *ηC*. The factor *η*, which describes the ratio of growth rates (i.e., *η* ≈ *r*_*C*_: *r*_*A*_) at low nutrient concentrations where the Monod growth is linear, is close to 1 due to the similarity in growth characteristics; see **Supplementary Note 1**.

**Figure N2.0:**
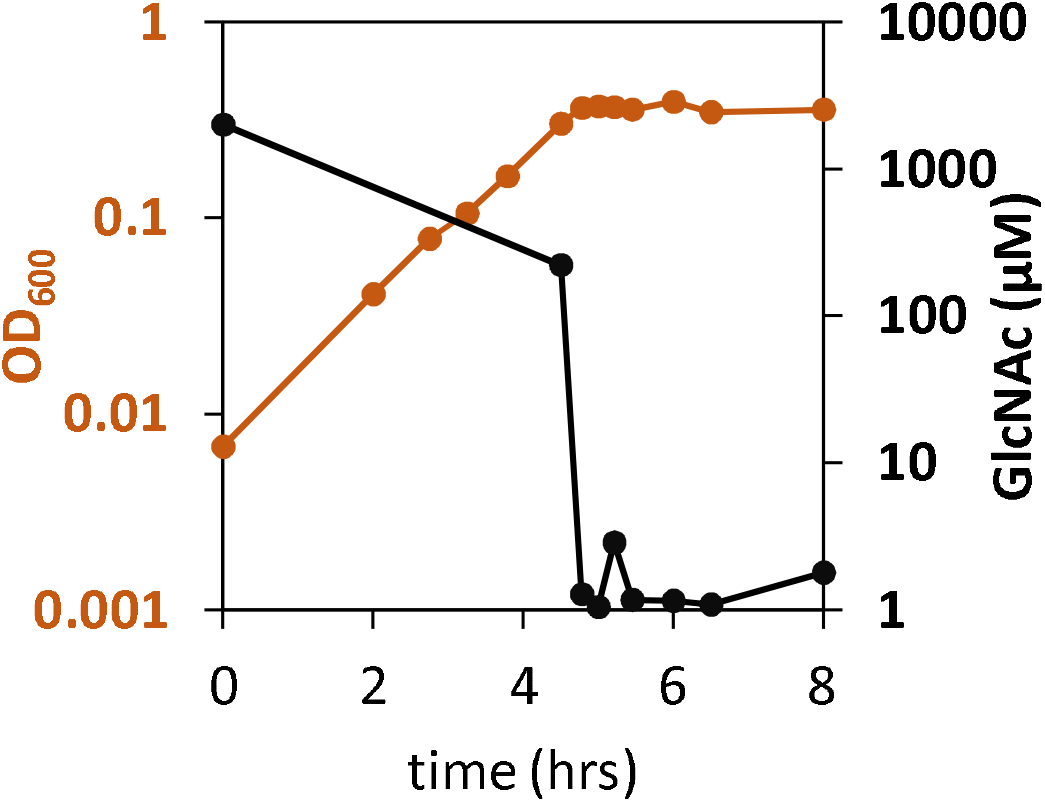
Growth of ViblAOl in GIcNAc batch culture. A batch culture of VTblAOl growing exponentially in GlcNAc is transferred to a fresh batch culture with 2mM GlcNAc at time 0 and monitored over the next 8 h. OD_600_ of this culture (orange) increased exponentially before halting abruptly at ∼0.3, where GlcNAc concentration (black) in the medium dropped to the order of 1 *μM*, and remained there the next several hours where the OD is saturated. The result indicates that the *K*_*M*_ of *Vib1AO1* for GIcNAc is in the range of 1*μM*. GIcNAc concentrations exceeding 250 μM, including those present before four hours, were above the limit of quantification and could not be accurately measured.

## 2. Cell growth and chitinase synthesis

We now apply the above considerations to the growth of chitin degraders and exploiters. The growth equations for the degraders and exploiters are respectively

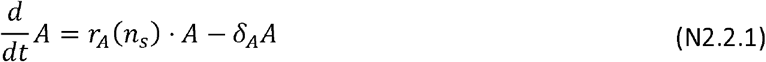

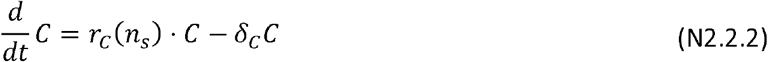

where the 2^nd^ terms on the right hand side of Eqs. (N2.2.1), (N2.2.2) describe cell detachment from chitin particles, with *δ*_*i*_ being the detachment rate of species *i*. The detachment of *Vib*AOl is well established experimentally^34^. A similar term is included for the exploiters. Also, we assume the detached cells do not find their way back to the chitin particles. [Reattachment was already small for chitin chips, and is expected to be even smaller for the much smaller colloidal chitin particles.]

Lastly, the equation describing the synthesis of chitinases by the degrader is

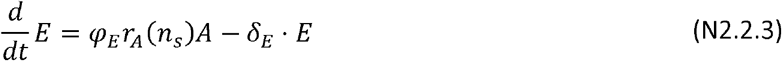

where we assume the chitinase production flux is a constant fraction *φ*_*E*_ of biomass production flux *r*_*A*_ · *A* (see Guessous et al.^34^), and *φ*_*E*_ describes the rate of chitinase loss (either detachment or turnover, also established in Guessous et al.^34^).

Using Eq. (N2.1.7) for R_*A*_(n_*s*_), Eqs. (N2.2.1)-(N2.2.3) become

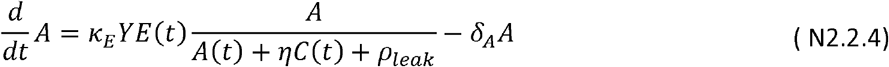

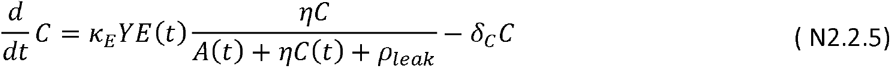

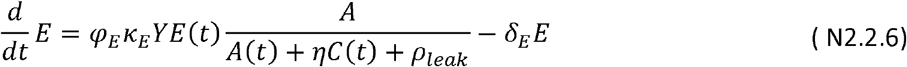

By defining an important effective parameter

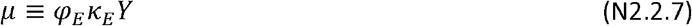

and re-expressing *E*(*t*) as *ε* (*t*) ≡ *E(t)/ φ*_*E*_, we obtain a more transparent set of equations:

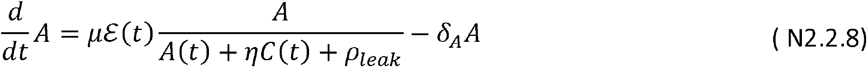

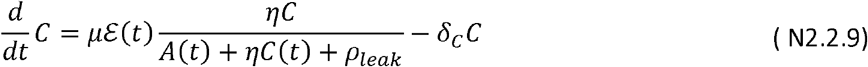

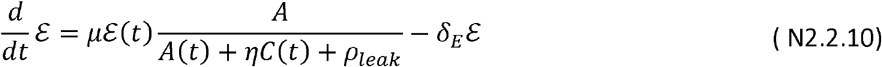

### 3. Analysis of systems dynamics

We first investigate a possible exponential growth phase admitted by Eqs. (N2.2.8)-(N2.2.10). From the forms of Eqs. (N2.2.8)-(N2.2.10), we see that if the density of the exploiters *C*(*t*) increases faster than that of the degraders, *A*(*t*), then the factor *A*/(*A* + *ηC* + *ρ*_*leak*_) will diminish over time and an exponential growth phase is not possible. We thus look for the exponential growth phase with 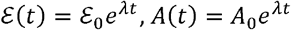, and 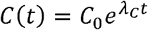, with the population growth rate for exploiters, *λ*_C_, being slower than that of the degraders, *λ*. Then Eqs. (N2.2.8)-(N2.2.10) lead to

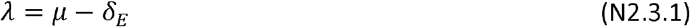

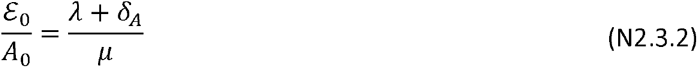

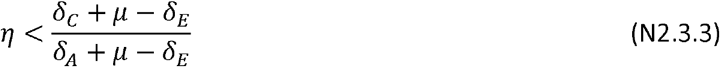

As *η ≈* 1 empirically (see Supp Note 1), condition (N2.3.3) is satisfied if the exploiters detach or turn over with a rate similar to that of the degraders. And given that the exploiter growth must be sub-leading in an exponentially growing culture, we shall focus on the dynamics of the degraders and the chitinases in the remainder, taking into account of the exploiters only through their effect on shifting *ρ*_*leak*_. Thus, we focus on the two-variable system

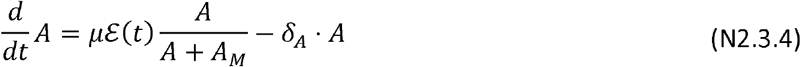

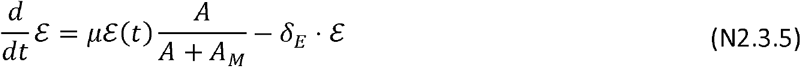

Where

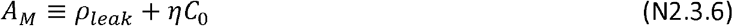

*C*_*0*_ being the initial exploiter density. To simplify the two-variable system further, we write Eqs. (N2.3.4) and (N2.3.5) in terms of the dimensionless variables, *a*(t) ≡ *A(t)/A*_*M*_ and *ε* (*t*) ≡ *ε (t)/A*_*M*_. *we* obtain the following pair of equations

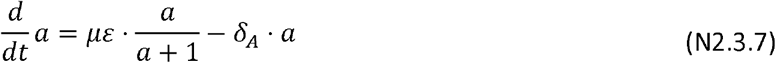

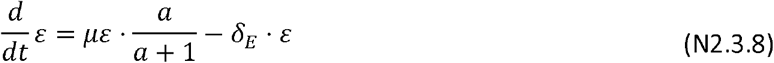

characterized by 3 parameters: *μ, δ*_*A*_, *δ*_*E*_

We next characterize the conditions that enable Eqs. (N2.3.7) and (N2.3.8) to admit the exponentially growing solution. First of all, we note that if the initial enzyme concentration is zero (i.e., *ε* (0) = *ε*_*0*_ = 0), then no growth is possible. This reflects the simplifying assumption we took in writing down Eq. (N2.2.3), that the chitinase production flux is proportional to the cell replication rate. Since cell replication depends on GlcNAc generated by chitinases, growth is not possible without chitinases to get the system started. In more realistic models, degraders must be able to synthesize some chitinases before generating substantial GlcNAc for uptake. This synthesis can be fueled by cellular carbon storage such as glycogen, converted from the degradation product of biomass components (e.g., the ribosomes, proteins, or lipids), or derived from the assimilation of residual GIcNAc monoers in the colloidal chitin culture. In this work, we do not model this initial dynamical phase, which requires deeper understanding of the cells’ physiology during starvation. Instead, we simply express the end result of this initial phase by an “initial” enzyme amount, *E*_*0*_ > 0, or *ε*_0_ > 0 in the dimensionless variable.

With *ε*_*0*_ *>* 0 (and a finite initial degrader concentration *a*_*0*_ *= a*(0) > 0), it is not always the case that the growth solution is obtained. From Eq. (N2.3.8), it is clear that no growth is possible if *μ < δ*_*E*_, i.e., the enzyme reproduction rate must exceed the enzyme loss rate. Restricting to *μ > δ*_*E*_, it is easy to see that if either *δA* or *δ*_*E*_ is zero, then exponential growth occurs for all *a*_*0*_ ≡ *a*(0) > 0 and *ε*_*0*_ ≡ *ε* (0) > 0: According to Eq. (N2.3.7), if *δ*_*A*_ *=* 0, then *a*(t) always increases for all *a*_*0*_ > 0 and *ε*_*0*_ *>* 0. As *a(t)* ≫1, *ε* (t) would grow exponentially according to Eq. (N2.3.8) as long as *μ > δ*_*E*_, and the exponential increase of in turn drives the exponential increase of. Alternatively, if, then will grow according to Eq. (N2.3.8) for all and; and when, would grow as well, resulting in the exponential growth of and hence. If both and, then the growth phase and is not always obtained for all initial conditions and. A separatrix separates the set of initial conditions that lead to exponential growth from those that do not, as shown by the red line in **Fig. N2.1**. For the monoculture (), crossing the red line is just the Allee transition.

**Figure N2.1:**
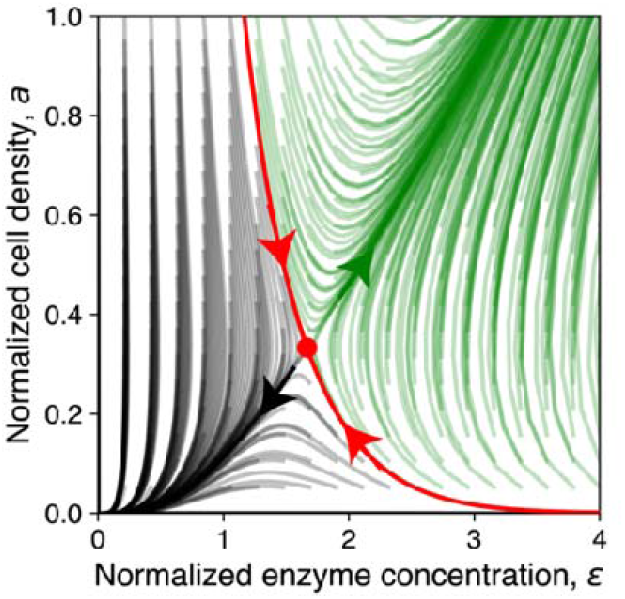
2-dimensional flow diagram generated by the two-variable dynamics Eqs. (N2.3.7) and (N2.3.8), showing how the cell and enzyme densities (and, respectively) evolve in time. Trajectories are color-coded based on whether they result in growth (green lines) or extinction (gray lines). The red line indicates the separatrix. The shape of this flow diagram only depends on two effective parameters, and which fix the position of the fixed point () indicated by the red circle. For this plot, we used and, using parameter values derived from **Table N2**.

The scale of the separatrix, i.e., the condition for which Allee transition occurs, is set by the values of the fixed point, and, at which and. We have and, which, in terms of the actual cell and chitinase densities, are:

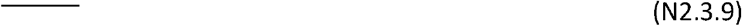

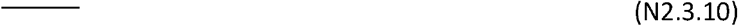

Thus, in order for this system to exhibit a transition between the growth and no growth phase, not only must there be the conditions, and, we must also have, i.e., nutrient loss (by diffusive leak or by stealing by exploiters) is *necessary* for a phase transition. Indeed, the scale of the phase transition as set by and is directly proportional to. For a system with large (such as the case for small colloidal chitin particles studied here), it is possible that the presence of a small amount of exploiters can shift the system from one side of the phase boundary to the other side, resulting in drastic change in the system’s fate, from growth to no growth.

### 4. Numerical solution

We obtained the separatrix by solving numerically Eqs. (N2.3.4), (N2.3.5) using the parameters listed in **Table N2**. The green line in **Fig. N2.2a** corresponds to the case of the degrader monoculture with no exploiters, and the grey line in **Fig. N2.2a** corresponds to a critical initial exploiter density, OD (or cells/ml) where the growth of the coculture ceased. To determine the operating point of the system, we note that the initial degrader density is OD (or cells/ml) as used in the experiments.

The initial amount of chitinases in the system,, which corresponds biologically to the amount of chitinases that the degraders produce during starvation, is not known. However, we can fix the value of within this model by requiring that it sits on the grey phase transition line. This is shown by the vertical dashed line in **Fig. N2.2a**, giving. Thus, we see the operating point is above the green separatrix, such that the system supports exponential growth in the absence of exploiters; but the operating point is on the grey separatrix, such that it does not grow exponentially in the presence of 0.001OD of exploiters. The actual trajectories in the phase space are shown as the dashed lines in **Fig. N2.2b** for (green), OD (red), and OD (grey).

**Figure N2.2:**
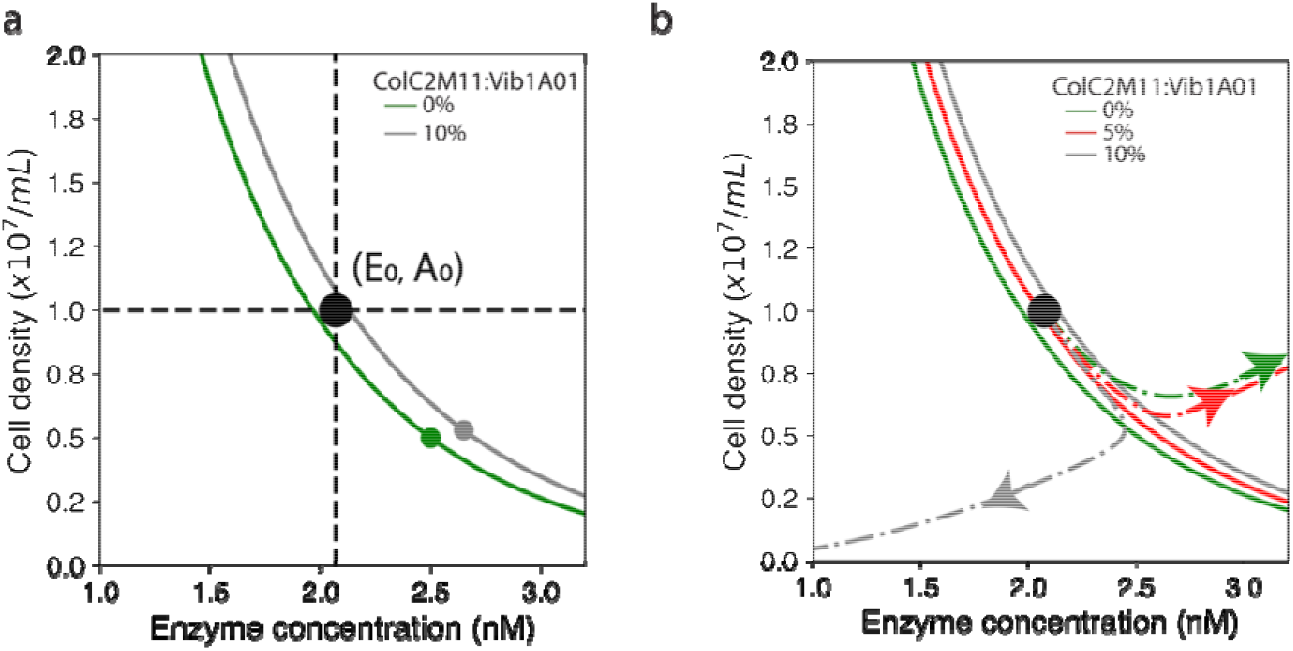
(a) Phase diagram in the initial cell and enzyme concentrations. The green line represents the separatrix for the monoculture () and the gray line represents the co-culture with. The solid green and gray circles represent the fixed points for these respective cases. The operational point is indicated at the crossing of the two dashed lines. (b) Trajectories in the A-E space for various initial exploiter densities are indicated as the dashed lines with arrows (colors indicated in the caption).

The time dependences of these 3 cases are shown in **Fig. 5E** of the main text. We find that for the intermediate case with the initial exploiter density being 5% of the initial degrader density (red line in **Fig. 5E**), the increase in lag over the monoculture lag is ∼20h, similar to observation (**Fig. 5F**). Overall, the lag time increases sharply as the critical point is approached (**Fig. N2.3**), demonstrating that this phase transition provides a qualitative metabolic-based mechanism, rationalizing the conundrum that a small increase in exploiter density destroys the growing phase.

**Figure N2.3:**
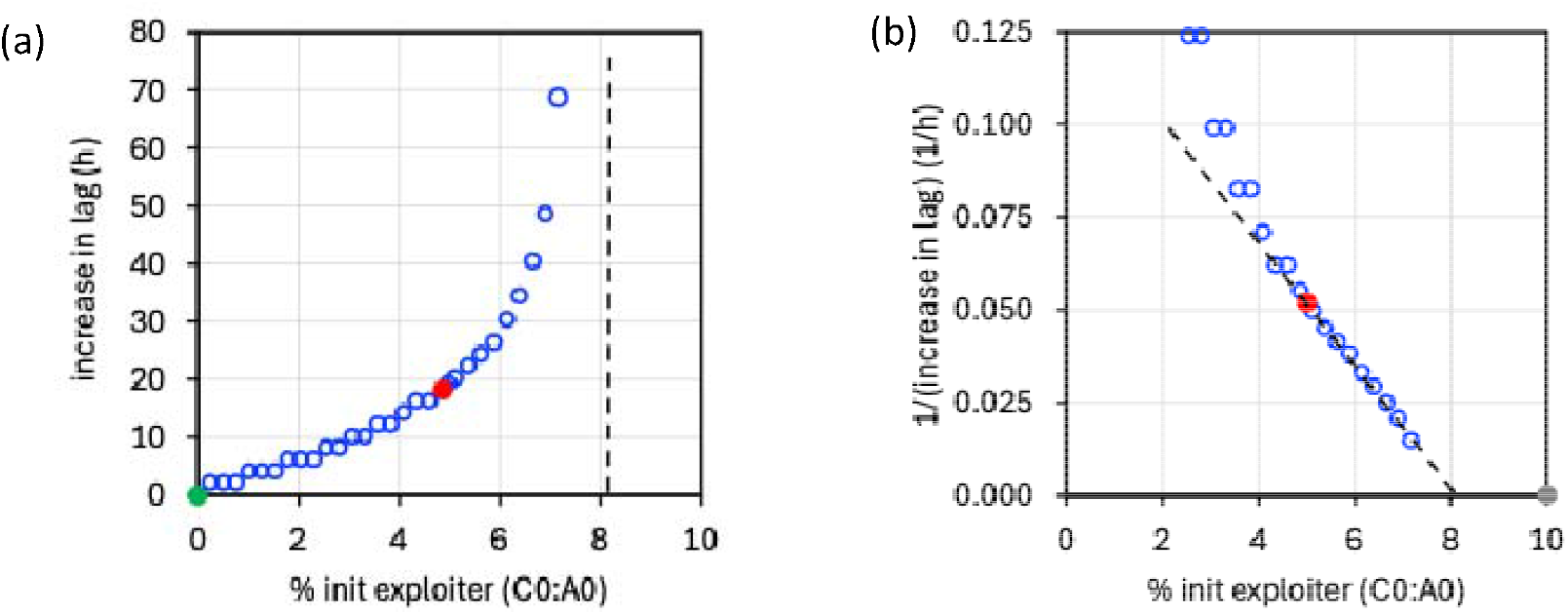
Divergence of lag time at phase transition. **(a**) Delay of the coculture compared to the monoculture for different initial values of the exploiter,. The green circle represents the monculture and the red circle the coculture with 5% *Col*C2M11 (see **Fig. 5E**,**F** of the main text). **(b)** Plotting the inverse of the increase in lag reveals the transition for this system occuring at *C*_0_ : *A*_0_ ≈ 8%where the increase in lag diverges. Dashed lines are drawn to guide the eyes.

However, the long lag time of the monoculture exhibited by the model (∼48h, **Fig. 5E**) is not consistent with the experimental observations (**Fig. 5F**). The long lag time reflects the difficulty of entering exponential growth. In our model, this may be a result of the simplifying approximation made in Sec. 1 of this Note, that bacteria covers the surface of the chitin particle uniformly. Such an assumption overlooks the possibility of higher initial bacterial densities on patches of a particle if bacteria are clustered closely together. A more sophisticated treatment would be required to provide a more realistic estimate of the lag time for degraders that start as a surface aggregate.

**Table N2:** Summary of parameters used in the numerical simulation. **a**, In this simulation study, cell density is measured in unit of OD_600_ which is taken to be a density of 10^9^ *cells/ mL* for both *Vib*1A01 and *Col*C2M11. **b**, The value of the Monod constant for *Col*C2M11 has not been measured and is not needed for the numerical solution. But its value can be inferred from the parameter *η*, whose value is given in the next row, together with the knowledge of the parameters *rmax, A, rmax,C*, and *K*_*A*_ listed in rows above. We find *K*_*C*_ ≈ 0.4 *μM*. **c**, The parameter *η*, which is the ratio of the growth rate of *Col*C2M11 and *Vib*1A01 at low GlcNAc concentrations, and is given by the ratio of the Monod parameters, is obtained from the result of fed-batch growth as described in **Supplementary Note 1**. **d**, The value of the catalytic rate of the chitinase, *K*_*E*_, was quantified in Guessous et al. for chitin chips. We expect it to be different (larger) for colloidal chitin studied here. The precise value of *K*_*E*_ is not needed for the model, which depends on *K*_*E*_ only through the lumped parameter *μ ≡ φ*_*E*_ *K*_*E*_ *Y*, whose value is provided in the next row. [From the values of, *μ φ*_*E*_, and *Y* provided, the value of *K*_*E*_ for colloidal chitin can be inferred and it is 2-3x larger than that found for chitin chips.] **e**, For the growing culture, the population growth rate *λ ≈* 0.16/*h* (**Fig. 5F**) is related to the chitinase regeneration rate *μ* and the chitinase turnover rate *δ*_*E*_ by Eq. (N2.3.1), *λ*= *μ* −*δ*_*E*_, which gives *μ ≈* 0.2/*h*. Note that the population growth rate *λ* is several times larger here for colloidal chitin compared to that obtained for chitin chips in Guessous et al. This is attributed molecularly to a larger chitinase activity *K*_*E*_ reflecting the ease of breaking down the chemically treated colloidal chitin.

| Parameter description | Symbol | Value | Unit | Reference |
| --- | --- | --- | --- | --- |
| GlcNAc diffusion coeff. | $D_n$ | 500 | $\mu m^2/s$ | |
| colloidal chitin radius | $R_0$ | 10 | $\mu m$ | Figure 4 |
| chitin volume fraction | $\phi_{chitin}$ | 0.14 | %v/v | Methods |
| nutrient leakage rate | $\gamma$ | 76 | $h^{-1}$ | Eq. (N2.1.6) |
| *Vib*1A01 cell density | $A$ | 0.01-1 | OD<sub>600</sub>; $10^9$ cell/ml | footnote a |
| yield on GlcNAc | $Y$ | 0.16 | OD<sub>600</sub>/mM GlcNAc | Refs. <sup>34,42,43</sup> |
| max *Vib*1A01 replication rate | $r_{max,A}$ | 0.6 | $h^{-1}$ | Figure 3A |
| max *Col*C2M11 replication rate | $r_{max,C}$ | 0.3 | $h^{-1}$ | Figure 3A |
| *Vib*1A01 Monod constant | $K_A$ | 1 | $\mu M$ | Ref <sup>34</sup>, Fig. N2.0 |
| *Col*C2M11 Monod constant | $K_C$ | 0.4 | $\mu M$ | footnote b |
| ratio of the growth rate of *Col*C2M11 and *Vib*1A01 at low GlcNAc concentrations | $\eta \equiv \frac{r_{max,C}/K_C}{r_{max,A}/K_A}$ | 1.2 | -- | footnote c, Supp Note 1 |
| population growth rate | $\lambda$ | 0.16 | $h^{-1}$ | Figure 5F |
| chitinase amount per total cellular proteins | $\varphi_E$ | 5% | -- | Ref. <sup>34</sup> |
| chitinase catalytic rate | $\kappa_E$ | | $\frac{mM \text{ GlcNAc}/h}{\text{enzyme}/mL}$ | footnote d |
| chitinase regen. rate | $\mu \equiv \varphi_E \kappa_E Y$ | 0.2 | $h^{-1}$ | footnote e<br>Eq. (N2.3.1) |
| *Vib*1A01 detachment rate | $\delta_A$ | 0.2 | $h^{-1}$ | Ref. <sup>34</sup> |
| chitinase turnover rate | $\delta_E$ | 0.04 | $h^{-1}$ | Ref. <sup>34</sup> |
| cell density whose uptake is equivalent to diffusive leakage flux | $\rho_{leak} \equiv \frac{\gamma K_A Y}{r_{max,A}}$ | 0.02 | OD<sub>600</sub> | Eq. (N2.1.8) |
| initial *Vib*1A01 density | $A_0$ | 0.01 | OD<sub>600</sub> | Figure 1D, 5F |
| initial *Col*C2M11 density | $C_0$ | 0-0.01 | OD<sub>600</sub> | Figure 1D, 5F |

## Footnotes

◊ The Monod constant *K*_*A*_ was indirectly estimated in Guessous et al^34^, and further confirmed here by direct measurement of the residual GIcNAc concentration for a ViblAOl monoculture grown in GIcNAc; see **Fig. N2.0**.

